# Structures of TGF-β with betaglycan and the signaling receptors reveal the mechanism whereby betaglycan potentiates receptor complex assembly and signaling

**DOI:** 10.1101/2024.07.19.604101

**Authors:** Łukasz Wieteska, Alexander B. Taylor, Emma Punch, Jonathan A. Coleman, Isabella O. Conway, Yeu-Farn Lin, Chang-Hyeock Byeon, Cynthia S. Hinck, Troy Krzysiak, Rieko Ishima, Fernando López-Casillas, Peter Cherepanov, Daniel J. Bernard, Caroline S. Hill, Andrew P. Hinck

**Affiliations:** Department of Structural Biology, University of Pittsburgh, Pittsburgh, PA, USA; Faculty of Biological Sciences, Astbury Centre for Structural Studies, University of Leeds, Leeds, UK; Department of Biochemistry & Structural Biology and Greehey Children’s Cancer Research Institute, The University of Texas Health Science Center at San Antonio, TX, USA; Chromatin Structure and Mobile DNA Laboratory, The Francis Crick Institute, London NW1 1AT, UK; Department of Pharmacology and Therapeutics, McGill University, Montreal, QC, Canada; Departmento de Biología Celular y del Desarrollo, Instituto de Fisiología Celular, UNAM, Mexico City, Mexico; Developmental Signalling Laboratory, The Francis Crick Institute, London NW1 1AT, UK

**Keywords:** TGF-β, Inhibin, co-receptor, TGF-β receptors, structural biology, betaglycan, NMR, X-ray crystallography, Cryo-EM, signal potentiation

## Abstract

Betaglycan (BG) is a transmembrane co-receptor of the transforming growth factor-β (TGF-β) family of signaling ligands. It is essential for embryonic development and tissue homeostasis and fertility in adults. It functions by enabling binding of the three TGF-β isoforms to their signaling receptors and is additionally required for inhibin A (InhA) activity. Despite its requirement for the functions of TGF-βs and InhA in vivo, structural information explaining BG ligand selectivity and its mechanism of action is lacking. Here, we determine the structure of TGF-β bound both to BG and the signaling receptors, TGFBR1 and TGFBR2. We identify key regions responsible for ligand engagement, which has revealed novel binding interfaces that differ from those described for the closely related co-receptor of the TGF-β family, endoglin, thus demonstrating remarkable evolutionary adaptation to enable ligand selectivity. Finally, we provide a structural explanation for the hand-off mechanism underlying TGF-β signal potentiation.

## Introduction

Secreted ligands of the Transforming Growth Factor-β (TGF-β) family transduce their signals by binding and bringing together type I and type II receptor serine/threonine kinases, which triggers a transphosphorylation cascade and activation of downstream signaling^1^. Similar to other cytokine families, some ligands of the TGF-β family bind co-receptors, which govern receptor binding and signaling^2^. One such co-receptor is the cell surface proteoglycan TGFBR3, also known as betaglycan (BG). Unlike the type I and type II receptors, BG lacks a cytosolic kinase domain and is not directly involved in transducing the signal^3^. BG is nonetheless essential in vivo, where it binds and potentiates the activity of the three TGF-β isoforms by presenting them to their signaling receptors, TGF-β receptor type I (TGFBR1) and TGF-β receptor type II (TGFBR2)^4^. Once the ligand–receptor signaling complex is formed in a stepwise manner, first by binding TGFBR2 and then by recruiting TGFBR1, TGFBR2 phosphorylates the co-complexed TGFBR1, which activates SMAD2 and SMAD3. TGFBR1 can also transphosphorylate other type I receptors, such as ACVR1(ref 5) or ACVRL1 (ref 5), activating the non-canonical SMAD1/5/9 pathway^6^.

BG binds the three TGF-β isoforms (TGF-β1, -β2 and -β3), 25 kDa disulfide-linked homodimers^7^ with comparable high affinity (K_D_ 5–20 nM)^8^. BG is most critical for potentiation of TGF-β2 signaling as this ligand binds TGFBR2 with 200–500-fold lower affinity than TGF-β1 and -β3. In the absence of BG, cells are largely unresponsive to TGF-β2, but in its presence, cells respond at concentrations comparable to the other isoforms^4, 9, 10^. BG null mice die during embryonic development and share many of the phenotypic characteristics of the TGF-β2 knockout, including severe heart and liver defects, consistent with the requirement of BG for TGF-β. BG is nonetheless important for the function of all three TGF-β isoforms, by sensitizing non-transformed cells to TGF-β mediated growth inhibition^9, 10^. Downregulation of BG by transcriptional repression or ectodomain shedding, a process by which BG is proteolytically cleaved and the soluble ectodomain is released into the extracellular space where it can bind and sequester TGF-βs from the signaling receptors^11^, correlates with oncogenic transformation in cancer cells^12–14^. Hence, BG is regarded as contributing to the tumor suppressive activity of the TGF-βs in vivo^11, 15, 16^.

In vivo, BG is required in gonadotrope cells for the activity of inhibin A (InhA), a TGF- β family heterodimer that blocks follicle stimulating hormone (FSH) secretion from the anterior pituitary^17^. InhA does this by forming a high-affinity complex that includes both BG and an activin type II receptor, ACVR2 or ACVR2B^18^, thereby sequestering the type II receptors from activins and other ligands which use them to activate signaling leading to expression of the FSH-β subunit^19^.

BG and a homologous co-receptor of the TGF-β family known as endoglin (ENG), which potentiates the function of BMP9 and BMP10, are transmembrane proteins with a large (approximately 760 amino acids) extracellular region comprised of two domains: the membrane-distal orphan domain and the juxtamembrane zona pellucida (ZP) domain^20, 21^. The orphan domain can be divided into two subdomains, OD1 and OD2, yet these are closely connected through paired antiparallel β-strands and adopt a well-defined orientation relative to one another^22^. The BG and ENG orphan domains (BG_O_ and ENG_O_, respectively) bind the TGF-βs and BMP9/10, respectively, though structural differences within the BMP9/10 binding interface between these domains suggest that TGF-β binding by BG_O_ is unlikely to mirror BMP9 binding by ENG_O22,23_. The ZP domain can also be subdivided into two subdomains, N-terminal ZP-N and C-terminal ZP-C. ZP-N is modified by two glycosaminoglycan (GAG) chains, though it does not contribute to ligand binding either directly or through its GAG chains in BG or ENG^24, 25^. The BG ZP-C domain (BG_ZP-C_) binds the three TGF-β isoforms and InhA^26^, while ENG ZP-C does not contribute to ligand binding and instead scaffolds the orphan domain for ligand binding. ZP-C is structurally similar to other ZP domains, including those in uromodulin, ZP3, and DMBT^27^. BG_ZP-C_ is also homologous to TGFBR3L, another co-receptor of the TGF-β family^28^ and serves to potentiate the inhibitory activity of inhibin B (InhB)^28, 29^. BG binds TGF-β homodimers with 1:1 stoichiometry, engaging with both the orphan and ZP-C domains^4, 8^. However, the isolated ZP-C domain binds TGF-β homodimers with 2:1 stoichiometry, effectively competing with TGFBR2 for binding^8^. In contrast, BG binds the α subunit of InhA via only its ZP-C domain^20, 30–32^ forming a 1:1 complex^8, 30^. Mutagenesis and NMR shift perturbation mapping of TGF-β and InhA are mostly in agreement, placing the interacting residues on the underside of the fingers, two β-hairpins that extend from the cystine knot core of each monomer^26, 33^. However, reports on how BG_ZP-C_ engages the ligands are contradictory^30, 34, 35^.

The uncertainty of how BG binds TGF-βs and InhA to potentiate their function is due to the absence of structural information for the complexes. This, together with the pivotal role that BG has on modulating the activities of TGF-βs and InhA, whose dysregulation can drive cancer progression^36^ and impact fertility^28^ respectively, motivated us to determine the structure of BG bound to TGF-β, which we have accomplished using X-ray crystallography and Cryo-EM. We also developed a new competition assay in living cells, which along with mutational analysis and predictions from Alphafold2 Multimer (AF2M)^37^, allowed us to identify the key determinants of ligand specificity. Together, this provides a detailed molecular-level understanding of how BG achieves ligand selection and potentiation of signaling, which involves sequential binding of the type II and type I signaling receptors, concomitant with stepwise displacement of the co-receptor^8, 38^.

## Results

### Structure of the TGF-β:BG complex

Initial trials to crystallize TGF-β2 bound to full-length BG complex presented difficulties, most likely due to the disordered 85-residue linker connecting the orphan and ZP domains. It was also not possible to determine the structure of this complex using Cryo-EM due to sample heterogeneity and strong preferential orientation. We therefore turned to a ‘divide-and-conquer’ approach and attempted to crystallize the TGF-β2:(BG_ZP-C_)_2_ and TGF-β2:BG_O_ complexes separately. The isolated TGF- β2:(BG_ZP-C_)_2_ complex again could not be crystalized, thus we turned to a previously engineered mini monomer of TGF-β2, mmTGF-β2 (ref 39). In spite of being a monomer and lacking the heel helix (Fig. 1A), mmTGF-β2 is much more soluble than TGF-β dimers and retains binding for BG^26^. This approach yielded high-quality orthorhombic crystals, enabling determination of the structure of the mmTGF-β2:BG_ZP-C_ complex at 1.9 Å resolution using X-ray crystallography (Fig. 1B, Supplementary Table 1).

**Figure 1.**
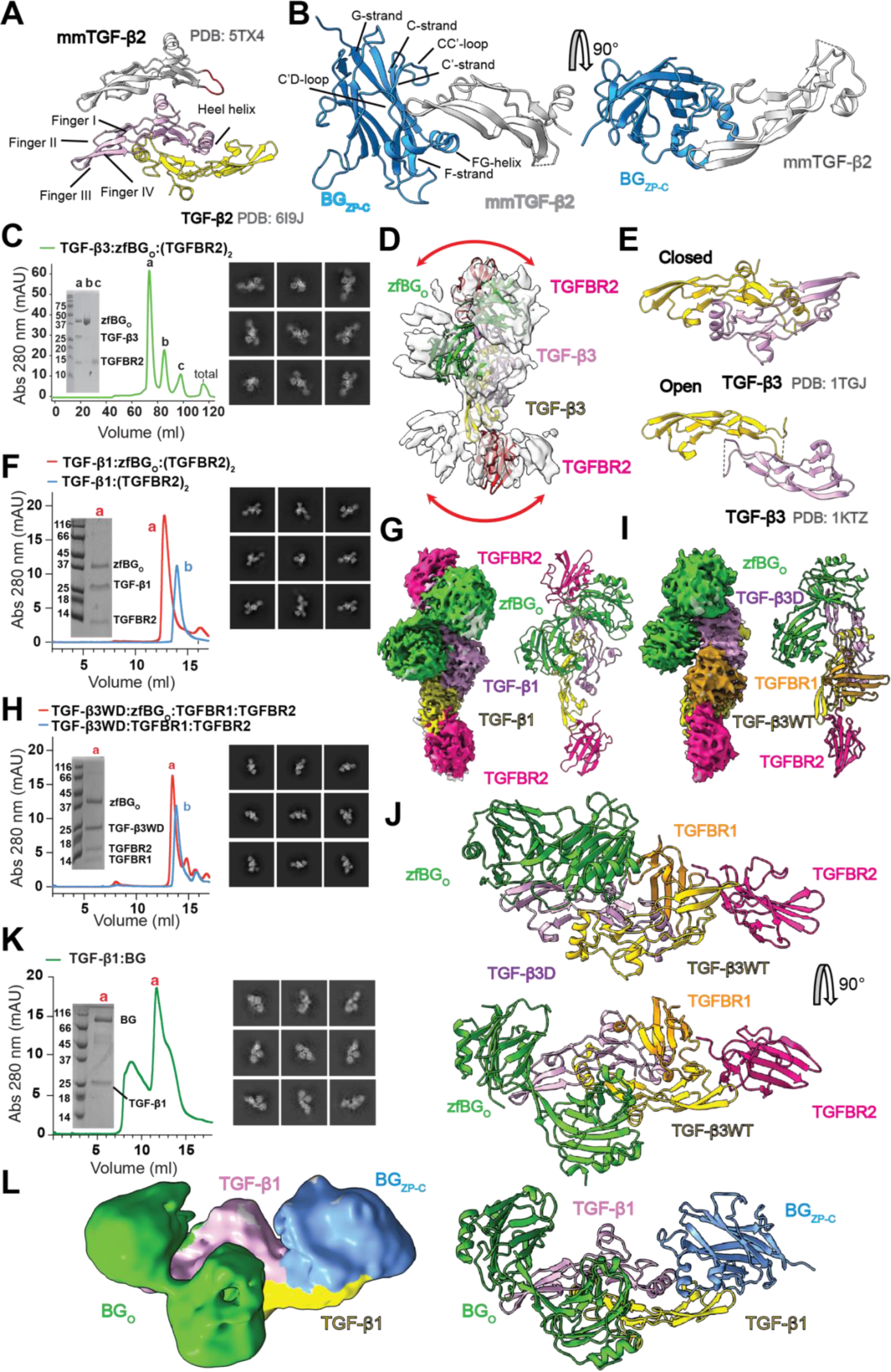
X-ray crystallographic and Cryo-EM structures of BG_ZP-C_ and zfBG_O_ in complex with TGF-β. (A) Structure comparison between TGF-β2 dimer and mmTGF-β2. The loop in the latter, which replaces the heel helix in the former, is highlighted in red. (B). Visualization of the X-ray crystallographic structure of mmTGF-β2 bound to BG_ZP-C_ with highlighted structural features of the ZP-C domain. (C) Size Exclusion Chromatography (SEC) purification profile demonstrating distinct peak of assembled TGF-β3:zfBG_O_:(TGFBR2)_2_ complex as assessed by SDS-PAGE (inset) and corresponding 2D classes. (D) Low resolution Cryo-EM map of the TGF-β3:zfBG_O_:(TGFBR2)_2_ complex. (E) Open-to-closed transition of TGF-β3 that is likely responsible for the low resolution of the Cryo-EM map. (F, H) SEC purification profiles demonstrating distinct peaks of assembled complexes of TGF-β1:zfBG_O_:(TGFBR2)_2_ (F) and TGF-β3WD:zfBG_O_:TGFBR2:TGFBR1 (H) as assessed by SDS-PAGE (inset) and corresponding 2D classes. (G, I) Respective Cryo-EM maps of imaged complexes (F, H respectively) with fitted structural models and highlighted, color-coded components. (J) Visualization of the X-ray crystallographic structure of the TGF- β3WD:zfBG_O_:TGFBR2:TGFBR1 complex. (K, L) SEC purification profile demonstrating distinct peak of assembled complexes of TGF-β1:ratBG as assessed by SDS-PAGE (inset) and corresponding 2D classes (K) with respective low resolution Cryo-EM map with highlighted components (L).

Unlike BG_ZP-C,_ BG_O_ does not prevent recruitment of TGFBR2 to TGF-βs, thus to understand the potentiation mechanism, we wanted to gain structural information on how the BG:TGF-β complex engages with the signaling receptors. We therefore attempted to crystallize the TGF-β3:BG_O_:(TGFBR2)_2_ complex. We used TGF-β3 instead of TGF-β2 to ensure high affinity binding of TGFBR2 and thus a stoichiometric complex. We also used zebrafish BG_O_, designated hereafter as zfBG_O_, rather than rat BG_O_ (ratBG_O_) owing to previous success crystallizing the former but not the latter^22^. While crystallization of this complex was not successful, we were able to obtain a low-resolution map of this complex using Cryo-EM and tentatively placed all components (Fig. 1C, D). We hypothesized that the expanded density in the areas attributed to TGFBR2 occurred due to a large-scale closed-to-open transition of TGF-β3 (Fig. 1E)^40^, which likely hindered high-resolution reconstruction and confounded efforts to obtain crystals.

We therefore sought to stabilize the TGF-β dimer in a single conformation and prepared an alternative complex replacing TGF-β3 with TGF-β1: TGF- β1:zfBG_O_:(TGFBR2)_2_ (Fig. 1F). We expected that the strong propensity of TGF-β1 to adopt the closed form^40^ would reduce or completely diminish the conformational heterogeneity. Additionally, in a separate approach we wanted to obtain the structure of the TGF-β:BG_O_ complex bound to both of the signaling receptors, TGFBR1 and TGFBR2, yet the TGF-β1:(TGFBR2)_2_:(TGFBR1)_2_ heterotetrameric assemble cannot recruit BG_O_, thus we aimed to construct a variant with the signaling receptors bound to only one monomer of TGF-β. We therefore switched to the TGF-β3 heterodimer, TGF-β3WD^41^, a mutated form that indeed can bind only one pair of TGFBR2:TGFBR1. We successfully assembled the TGF-β3WD:zfBG_O_:TGFBR2:TGFBR1 complex (Fig. 1H) and in this case we used the more flexible TGF-β3, expecting that the bound TGFBR1, which spans the TGF-β dimer interface, would lock TGF-β3 in the closed conformation^42^. Cryo-EM analysis of both of these complexes yielded density that enabled domain placement without ambiguity and closely accommodated the component structures (Fig. 1G, I and Supplementary Table 2). Despite this, it was difficult to build molecular models due to strong preferential alignment of the particles (Extended Data Fig. 1A, B). Thus, we returned to crystallography and successfully obtained orthorhombic crystals of the TGF-β3WD:zfBG_O_:TGFBR2:TGFBR1 complex and determined the structure at a resolution of 3.0 Å (Fig. 1J, Supplementary Table 1).

With knowledge of the substructures of the two component domains of BG bound to TGF-β, we repeated the analysis of full-length rat BG in complex with TGF-β, using TGF-β1 instead of TGF-β2. We successfully derived a Cryo-EM map of the entire complex, albeit at low resolution, into which we were able to unambiguously position the BG_ZP-C_ and BG_O_ domains onto the TGF-β dimer, which agreed with the positioning obtained from the substructures (Fig. 1L, Extended Data Fig.1F–G, Supplementary Table 2). Further, the full-length structure revealed no additional density corresponding to BG_ZP-N_, suggesting that the positioning of this domain is not fixed relative to the BG_ZP-C_. We also confirmed that BG_O_ and BG_ZP-C_ are placed on the opposite monomers of TGF-β, without any contact between domains (Fig 1L, Extended Data Fig. 1G).

In addition to our experimental pursuits, we explored the potential of AF2M^37^ to predict the structures of these complexes. While AF2M capabilities were challenged when full-length BG complexed with TGF-β2 was modeled, the predictions were impressively accurate for the individual domains, BG_O_ and BG_ZP-C_ complexed with TGF-β2 (Extended Data Fig. 1H, I). This demonstrates the utility of this recent technology for simulating the protein–protein interactions we aim to visualize. We have leveraged this capability to gain a better understanding of substrate specificity, as described below.

### Betaglycan ligand specificity

Among the 33 members of the TGF-β family, BG is known to bind with high affinity to the three TGF-β isoforms through both the BG_ZP-C_ and BG_O_ domains^20, 43^ and to the α subunit of InhA through the BG_ZP-C_ domain^28^. It was proposed that BG can also bind to various BMPs^44, 45^, but biophysical measurements contradicted these claims^26^. To determine the BG ligand selectivity, we developed a competition assay to test if selected ligands of the TGF-β family can compete with TGF-β2 for binding BG. For this, we established a HEK293T cell line stably expressing rat BG with an N-terminally attached SNAP-tag (Extended Data Fig. 2A-B) that allows for specific labelling through click chemistry^46^. After confirming that the SNAP tag did not negatively impact BG activity (Extended Data Fig. 2C) and that fluorescently labeled SNAP-BG signal colocalizes with fluorescently labeled TGF-β2 (Fig. 2A and Extended Data Fig. 2D), we assessed the ability of 100-fold excess of unlabeled ligand to compete with the labeled TGF-β2. Labeled TGF-β2 bound to SNAP-BG could be efficiently outcompeted by unlabeled TGF-β1, TGF-β2 and TGF-β3, however ActA, BMP2 and BMP4 were not able to compete (Fig. 2A, B). We further demonstrated that the co- localization of fluorescently labeled BG and TGF-β could be decreased by unlabeled TGF-βs in a dose-dependent manner (Extended Data Fig. 2D). Together, these data suggest that among the ligands tested, only TGF-β isoforms can bind BG with high affinity.

**Figure 2.**
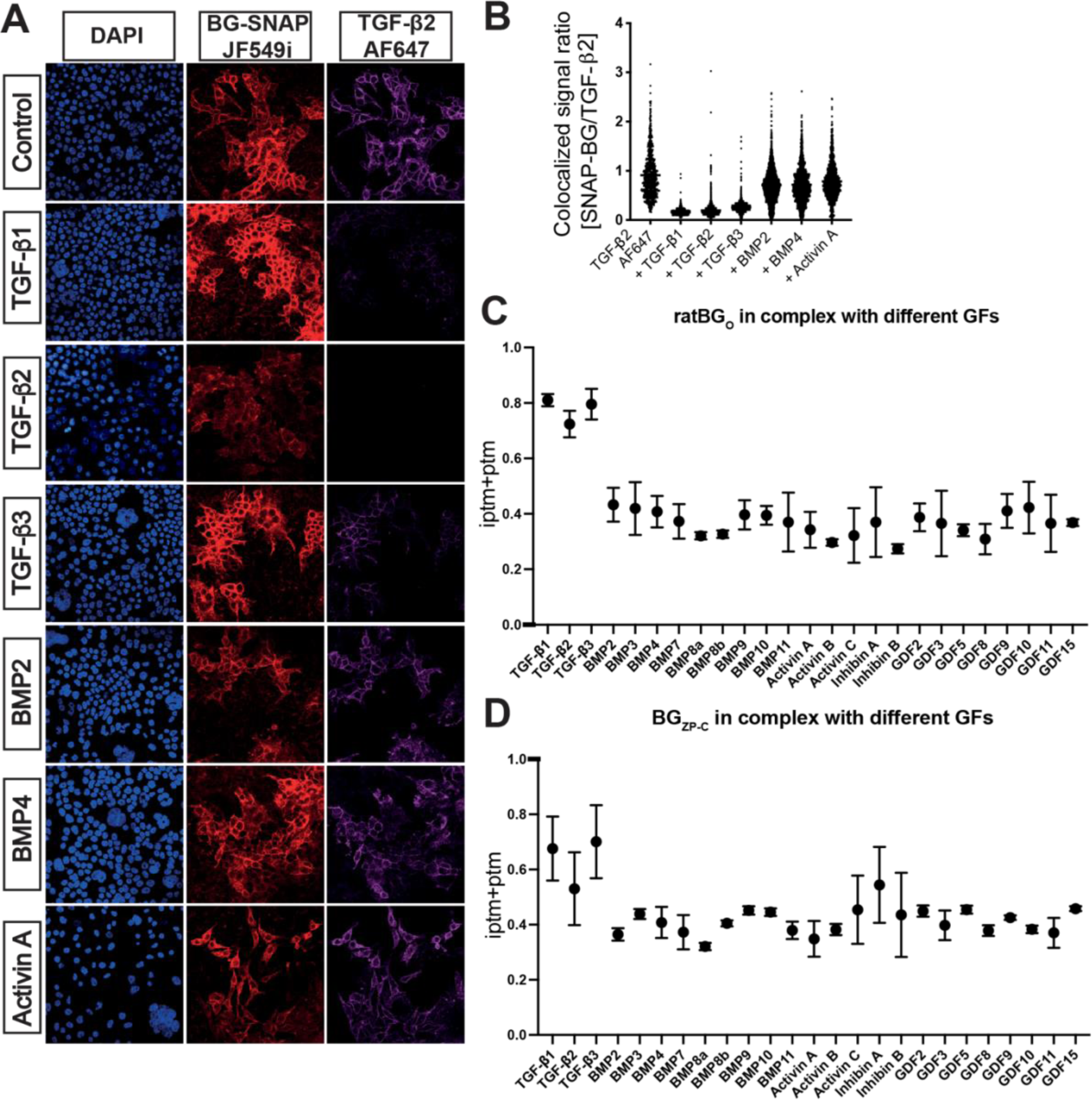
Selectivity of BG towards TGF-β family ligands. (A) Confocal microscopy images of HEK293T cells stably transfected with a SNAP-tagged BG construct (SNAP-BG) stained with DAPI (left panels, blue), incubated with SNAP-tag ligand conjugated to the CF567 fluorescent dye (middle panels, red), along with TGF-β2 tagged with AF647 fluorescent dye (right panels, violet) alone (row 1) or with the addition of the indicated unlabeled ligands (rows 2–7) (B) Quantification of the fluorescent colocalized SNAP-BG/TGF-β2 microscopy data. (C, D) AF2M analysis for ratBG_O_ (C) and BG_ZP-C_ (D) complex formation capability with selected ligands. The Y-axis represents the iptm+ptm score, which is a predictive measure of protein–protein interaction strength based on structural modelling.

Building upon the success of our structural predictions using AF2M, we expanded our survey by integrating AF2M modelling to encompass a broader spectrum of growth factors within the TGF-β family. The ptm+iptm score computed for BG_O_ or BG_ZP-C_ in complex with the TGF-βs, which provides a probability estimate assessing both post-translational modifications (PTMs) and inter-residue contacts (iPTMs) within the protein structures, was significantly higher than for other ligands, suggesting that BG is much more likely to bind TGF-β isoforms compared to any other TGF-β family ligand (Fig. 2C, D), which agrees well with our competition assay. As expected, predictions for InhA binding to BG_ZP-C_ also produced an elevated ptm+iptm score compared to other ligands, in agreement with the experimental data (see Fig. 6). In addition, the lower score for InhA compared to TGF-β isoforms, likely reflects lower binding affinity compared to TGF-βs^28^. However, it should be noted that the AF2M predictions for BG_ZP-C_ span a narrower range between true positive and negative cases compared with those for BG_O_, suggesting it is important to be cautious in interpreting the BG_ZP-C_ results.

### Interactions between TGF-β and the ZP-C domain

The structure presented in Fig. 1B shows that BG_ZP-C_ uses the surface of the exposed sheet formed by the C, C’, F, and G strands to pack against the fingertips and the inner surface of fingers I-II and III-IV of mmTGF-β2. The BG_ZP-C_ binding site on TGF-β lies on the inner surface of the fingers of TGF-β2, involving residues Leu^28^, Ile^33^, Val^61^, Leu^86^, Ile-^88^, Ile^92^, Lys^97^, Glu^99^, and Leu^101^. It was previously suggested that Ile^92^, Lys^97^, and Glu^99^, which are essential for binding BG_ZP-C_, are conserved among the TGF-β isoforms and inhibin α subunit, setting these ligands apart from other members of the family^26^. In the crystal structure of the mmTGF-β2:BG_ZP-C_ complex, Val^92^, Lys^97^, and Glu^99^, which lie within fingers III and IV, indeed form key contacts with BG_ZP-C_ (Fig. 3A I–III). In addition, the concave surface of TGF-β finger IV forms hydrophilic interactions with the irregularly structured region of the FG loop, particularly mmTGF-β2 Lys^97^, Tyr^90^, Glu^99^, and Lys^97^ which interact with BG_ZP-C_ Leu^156^, Ala^158^, Thr^159^, and Leu^156^, respectively. It is notable that most of these interactions are not formed between side chains, but rather between the side chain of one residue and the backbone amide or carbonyl of another residue. The tips of the fingers of the ligand are also engaged and form contacts with residues located on the CC’- (Pro^71^, Asn^72^) and C’D-loop (Asn^82^, Lys^96^, Asp^88^, as well as C’ strand (Glu^81^), with Asn^82^ being vital as assessed by Surface Plasmon Resonance (SPR) (Fig. 3B, C).

**Figure 3.**
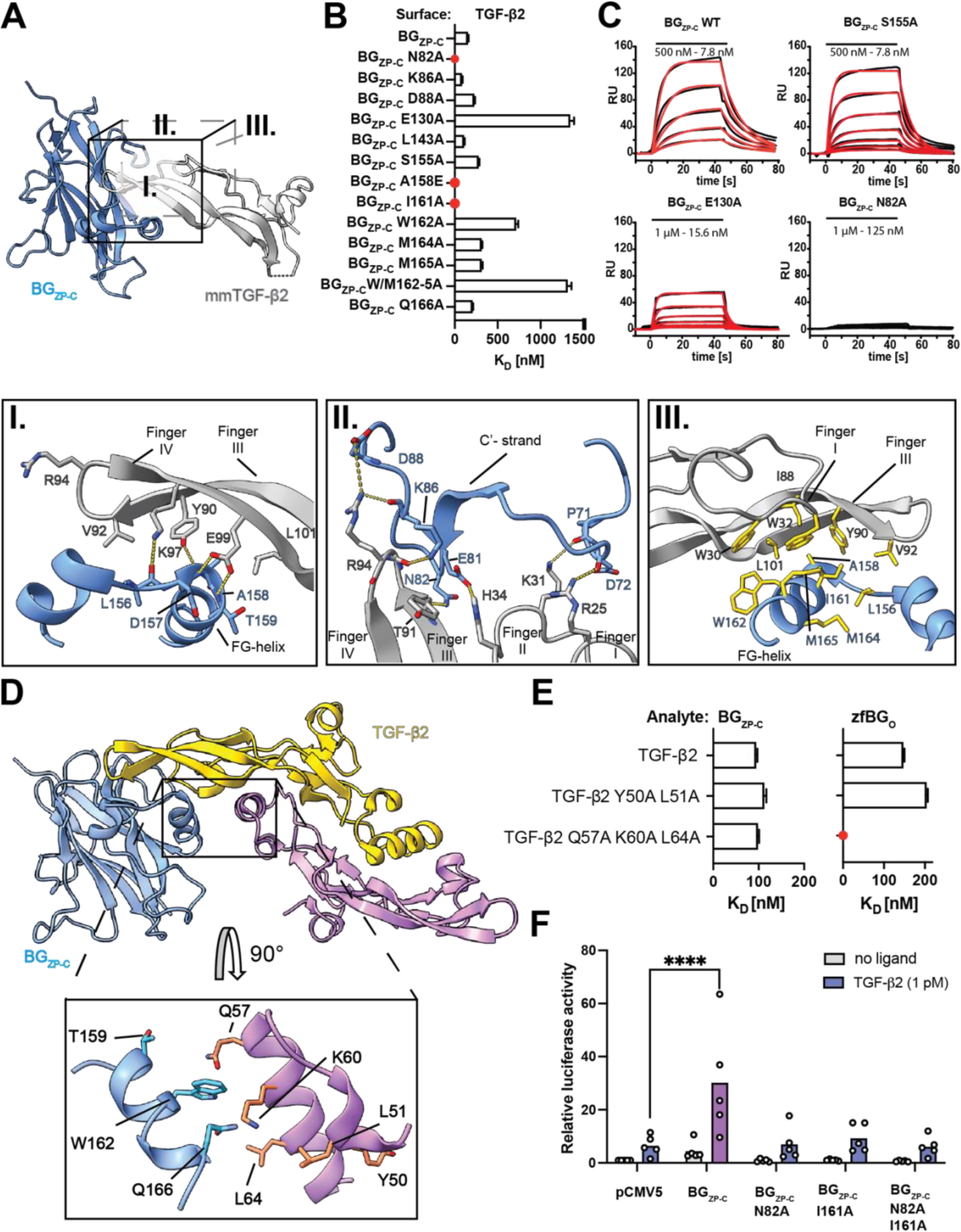
The TGF-β2:BG_ZP-C_ interface. (A) Visualization of the binding interface between BG_ZP-C_ and TGF-β2. Detailes of the interface are shown magnified below. The orientations are denoted by the sides of the cube. (I) Hydrophilic interactions between the extended region of finger I and the FG- loop of BG_ZP-C_, and between the fingertips and CC’-loop and C’-strand (II) are highlighted and denoted by dashed lines representing intermolecular bonds. (III) Hydrophobic contacts form between the concave surface of the fingers and the FG- helix that forms upon growth factor binding. Side chains of residues contributing to the hydrophobic interactions are highlighted in yellow. (B) Chart summarizing SPR data of BG_ZP- C_ mutant variants. Mutations N82A, I161A, and A158E in BG_ZP-C_ that completely abrogate binding to mmTGF-β2 are highlighted in red. (C) Selected SPR binding sensorgrams of BG_ZP-C_ mutants binding to immobilized TGF-β2 on the chip surface. (D) AF2M model of TGF-β2 in complex with BG_ZP-C_,with blowup panel highlighting potential interaction between BG_ZP-C_ FG- and TGF-β2 heel helices. (E) Chart summarizing SPR binding data on selected combined mutants (Q57A, K60A, L64A of the heel helix and Y50A, L51A of the pre-helix extension) binding to BG_ZP-C_ and zfBG_O_. (F) The mutants that most strongly perturbed binding in (B) were further investigated using a CAGA_12_-luciferase (CAGA-luc) assay for signaling activity in L6E9 cells.

In earlier structures of rat and mouse BG_ZP-C_ domain alone, the FG loop, which bridges the penultimate and ultimate β-strands (F and G, respectively), had weak electron density and was either modeled in an extended geometry^34^ or not at all^35^. In contrast, in the structure of the mmTGF-β2:BG_ZP-C_ complex, the FG loop undergoes a disorder-to-order transition and forms a helix that also packs against the inner surface of fingers I-II and III-IV of mmTGF-β2 (Fig. 3A III). Hydrophobic residues Ala^158^, Ile^161^, Trp^162^, and Met^165^ form crucial interactions with the concave surface of the growth factor fingers, including Trp^30^, Trp^32^, Val^92^ and Leu^101^ (Fig. 3A III) and mutations within this region severely disrupted binding (Fig. 3B).

It was reported that the affinity of mmTGF-β2 for binding BG_ZP-C_ is weakened by 10- fold compared to TGF-β2 (ref 26). However, an overlay of mmTGF-β2 with a full TGF- β2 monomer does not suggest any additional contacts that could account for the affinity difference. Hence, we considered the possibility of additional interactions between BG_ZP-C_ and the opposing monomer of TGF-β that are absent in mmTGF- β2:BG_ZP-C_. Indeed, alignment of the mmTGF-β2 as bound to BG_ZP-C_ onto the TGF-β2 homodimer suggested that the FG-helix and heel helix/pre-helix extension of the second monomer might contact one another. This extends the interface in this area and is consistent with the positioning of BG_ZP-C_ in the TGF-β1:BG structure (Fig. 1L) and an AF2M model of the TGF-β2:BG_ZP-C_ complex (Fig. 3D). To investigate this, we simultaneously mutated Tyr^50^, Leu^51^ and Gln^57^ or Lys^60^ and Leu^64^ in the pre-helix extension or helix region of TGF-β2 where the conserved residues could engage BG_ZP-C_. However, both mutants displayed only marginally reduced binding affinity towards the BG_ZP-C_, indicating other residues contribute to binding, or additional interactions are formed between side chains and the backbone in the proposed interface (Fig. 3E and Extended Data Fig. 3A).

To investigate the functional impact on TGF-β signaling potentiation of the key residues identified above, we co-transfected L6E9 rat muscle myoblasts, which express little to no BG, with BG_ZP-C_ constructs with Asn^82^ or Ile^161^ substituted with Ala, alongside a TGF-β responsive CAGA_12_-luciferase reporter construct^47^. Upon stimulation of the transfected cells with low concentrations of TGF-β2, mutation of both Asn^82^ and Ile^161^ markedly diminished CAGA_12_-luciferase activity in comparison to wildtype (WT) BG_ZP-C_ (Fig. 3F). These findings confirm that binding-deficient BG_ZP-C_ mutants diminish TGF-β signal potentiation.

To further validate that the BG_ZP-C_ mutations primarily influence ligand binding, without disrupting the structure, we recorded NMR natural abundance 2D ^1^H-^13^C shift correlation and 1D ^1^H spectra focused on the amide region of selected mutants (Extended Data Fig. 3B–E). Comparison of the spectral fingerprints of the mutant BG_ZP-C_ proteins against those of WT BG_ZP-C_ confirmed that the mutations did not disrupt the structures. This supports the conclusion that Asn^82^ and Ile^161^ play critical roles in TGF-β signaling by specifically binding TGF-β ligands.

### Interaction between TGF-β and the orphan domain

It was not known how the BG orphan domain engages the TGF-βs. Initial assumptions were drawn from the structure of the closest homolog, ENG, bound to BMP9 (ref 23). However, our structural data demonstrates a completely novel interface, with the zfBG_O_ subdomains, OD1 and OD2, and the paired antiparallel β strands that tether them together, forming an arc that straddles the edge of the TGF-β fingers (Fig. 4A and Fig. 1J). The positioning of zfBG_O_ on TGF-β is responsible for the 1:1 stoichiometry as a significant clash between OD1 from both protomers would occur with a 2:1 stoichiometry (as in case of ENG_O_) (Fig. 4B). The most extensive and crucial interface for TGF-β binding is formed by the loop from residues 244–251 located between the β2 and β3 strands of OD1 with the sequence PNSSSAFQ. The binding loop (BL) lies on the concave surface of the fingers and extends to the heel helix of the neighboring monomer (Fig. 4A I). The replacement of BL with an unrelated sequence of similar length (ΔBL) resulted in complete loss of binding (Extended Data Fig. 4C). Residues Asn^245^, Ser^247^, and Gln^251^ form hydrophilic interactions with finger I and II of TGF-β, while Ala^249^ and Phe^250^ engage Trp^30^, Trp^32^, Leu^86^, Leu^101^, Met^104^ within the concave surface (Fig. 4A I) of the fingers and Thr^57^ in the proximal part of the heel helix of the opposing monomer (Fig. 4A III). Mutations in this region, such as F250A or N253A in zfBG_O_ (Fig. 4C and Extended Data Fig. 4A), or in the heel helix of TGF-β that is contacted by the BL, such as the Q57A, K60A, L64A triple mutant of TGF-β2 described earlier (Fig. 3E and Extended Data Fig. 3A), either eliminated or significantly impaired binding. One caveat to these results is that the N253A mutation appeared to perturb the structure, as revealed by 1D ^1^H NMR, though other substitutions in zfBG_O_ did not (Extended Data Fig. 4B). On the other subdomain, OD2, we observed that residues Leu^208^, Leu^210^, and Leu^213^, which are located on the loop between β10 and the OD2 exit strand, pack against the outer convex surface of the fingers (Fig. 4A II). Mutation of any of these residues to Ala also dramatically reduced binding, with the latter two eliminating binding completely (Fig. 4C and Extended Data Fig. 4A). The interface is closed with Phe^49^ located at the end of the linker β-strand, mutation of which was also detrimental to ligand binding (Fig. 4C).

**Figure 4.**
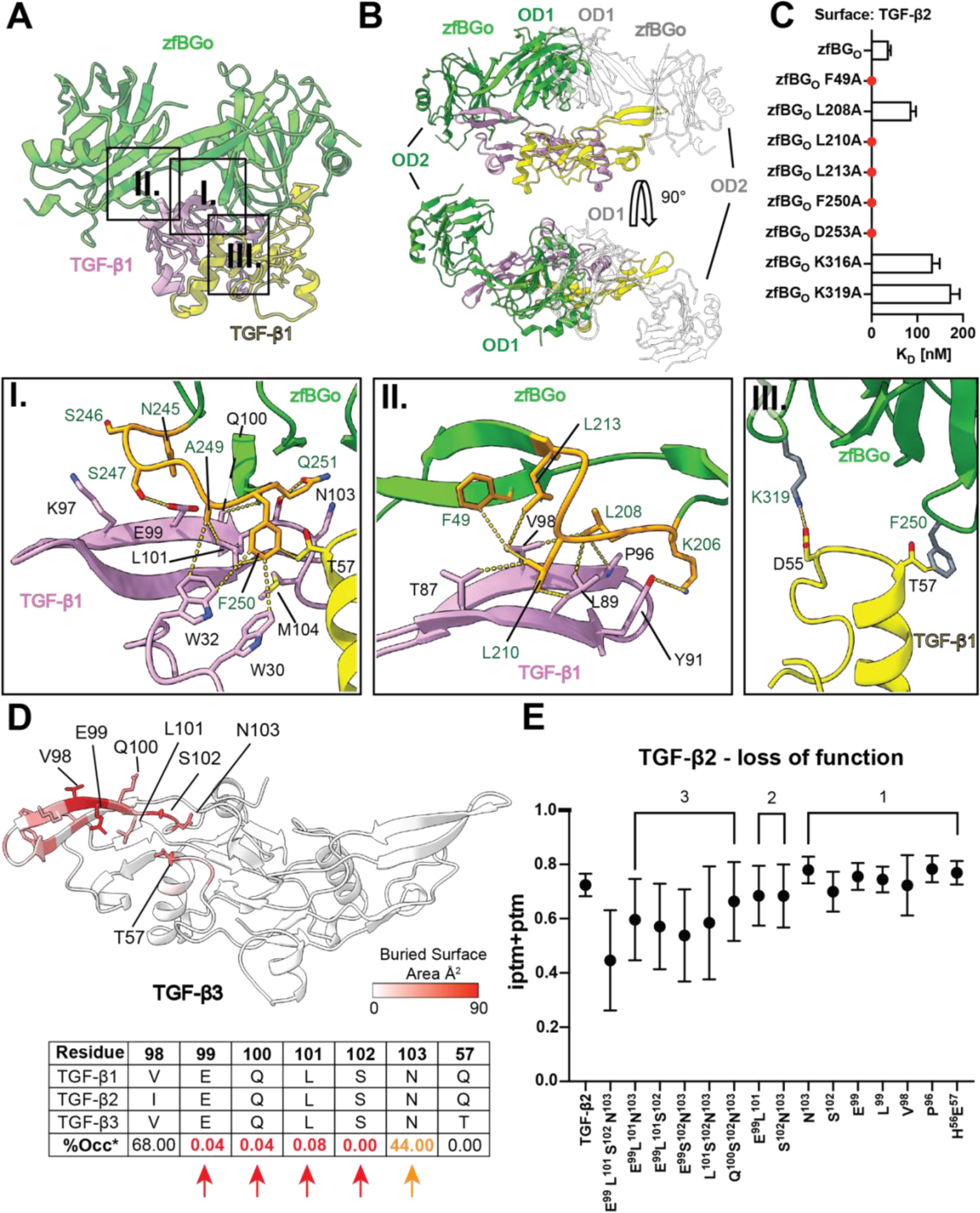
The TGF-β:zfBG_O_ interface. (A) The TGF-β:zfBGo interface (regions highlighted enlardeg below) is mostly maintained through hydrophobic interactions (I, II) between the ligand fingers and the binding loop (BL) present on OD1 (II) and a second, shorter adjacent loop present on OD2 (I). (III) Interactions between zfBG_O_ and the heel helix of the second monomer of TGF-β. (B) Conformation of zfBGo bound to TGF-β explains the 1:1 stochiometry as the second protomer of zfBG_O_ (white) cannot occupy binding area on the second monomer of TGF-β (yellow) due to significant structural clash between OD1 subdomains of both protomers. (C) Chart summarizing SPR binding data on selected interface mutants of zfBG_O_. (D) Structure of TGF-β3 coloured after Buried Surface Area (BSA) demonstrating contribution of each residue into the binding interface when in complex with zfBG_O_. The calculation was performed using the PISAePDB webserver. Below, table showing residue conservation among TGF-β isoforms and % of the occurrence of that residue in other Growth Factors from TGF-β family (E) Selected residues in TGF-β2 based on PISA calculations were mutated *in-silico* into the equivalent BMP2 residues to assess their contribution to binding by AF2M. Single, double, triple and quadruple combinations were tested, showing their additive effect and the most disruptive combination.

The identification of critical residues in the zfBG_O_ interface paved the way for a deeper understanding of specificity of zfBG_O_ towards TGF-β family ligands. Inspection of the interface buried area and TGF-β family Multiple Sequence Alignment (Fig. 4C, D), led to recognition of potential residues characteristic to only TGF-β isoforms, namely Glu^99^, Leu^101^, Ser^102^, or Asn^103^. The Gln^57^ to Ala mutation in the heel helix of the adjacent monomer was probably the key residue in triple mutant that impacted BG_O_ binding (Fig. 3E). However, since this residue is not conserved among TGF-β isoforms, most likely the proper positioning of the heel helix is essential. The AF2M simulations indicated that while substitution of each of these residues with the equivalent residue in BMP2 only moderately diminished zfBG_O_ binding, combinations of these amplified disruptive effect, with the quadruple variant of Glu^99^, Leu^101^, Ser^102^, and Asn^103^ placing the complex prediction score among the non-binders (Fig. 4E). Unfortunately, it was not possible to evaluate the effect of these substitutions *in vitro*, as this quadruple variant and related triple or double variants could not be isolated, suggesting that these residues must be playing further structural roles.

### The mechanism behind signal potentiation

Co-receptors, while not capable of signaling on their own, enhance the signaling potency of ligands via their primary receptors, which has important implications for the precise modulation of cellular signaling^3,4, 16^. There are several mechanisms through which co-receptors can act, including increasing the effective ligand concentration at the plasma membrane^23, 48, 49^, potentiation of signaling receptor binding by allosteric modulation^50^ or by direct contact with the signaling receptor^51^, or mediation of receptor internalization^52^.

Signal potentiation by ENG is thought to rely on efficient capture of BMP9/10 on the cell surface, increasing the local concentration and propensity of the ligands to bind and recruit their type I receptor, ACVRL1 (ref 23). It was suggested that BG not only potentiates signaling using the same mechanism as ENG, but also by indirectly or directly increasing the binding affinity between TGF-βs and TGFBR2 through its orphan domain^8^. Our structure of BG bound to TGF-β supports this dual mechanism as BG embraces the ligand homodimer in a way that allows one molecule of TGFBR2 to bind, thus allowing for the signaling receptor recruitment to the complex by simply “presenting” the ligand to the signaling receptor. In addition, the proximity of bound BG_O_ to recruited TGFBR2 suggests possible contact between the co-receptor and receptor (Fig. 1G) and thus direct potentiation. To confirm the direct potentiation mechanism, we conducted an SPR potentiation assay using both zfBG_O_ and ratBG_O_. It was surprising to discover that in contrast to ratBG_O_, zfBG_O_, which we used for X- ray crystallogaphy, did not potentiate the binding (Fig. 5A–C). ratBG_O_ and zfBG_O_ share 55.4% sequence identity, with no obvious difference that we could identify to explain the observed effect. To decipher the direct potentiation mechanism, we analyzed the TGF-β1:BG_O_:(TGFBR2)_2_ complex by Cryo-EM where zfBG_O_ was replaced with ratBG_O_ (Extended Data Fig. 5A–C). The strong preferential orientation observed previously was overcome using graphene oxide grids (Extended Data Fig. 5D) enabling building of a model into the 3.48 Å resolution Cryo-EM map (Supplementary Table 2). From the structure it was apparent that the most pronounced difference lay within the TGF- β binding loop of OD1, where there is a two-residue insertion in ratBG_O_ compared to zfBG_O_ (Fig. 5D, E) and that extension of this loop allows the BL in ratBG_O_ to make additional contacts with TGFBR2. To test our model, we swapped the shortened loop from zfBG_O_ into ratBG_O_. Despite similar binding affinity for TGF-β2 (Extended Data Fig. 5E and Supplementary Table 3), potentiation of TGFBR2 binding was not detected (Fig 5F, H), showing that the extension in the binding loop is essential for potentiation. In the structure of the TGF-β2:ratBG_O_:(TGFBR2)_2_ complex, Tyr^248^ is predicted to be responsible for most of the contact with TGFBR2 Phe^10^, Pro^11^, Leu^13^ and Ile^39^ (Fig. 5E). Indeed, mutation of ratBG_O_ Tyr^248^ to Ala nearly eliminated the potentiation effect, in accordance with our model (Fig. 5G, H). This unexpected difference between zfBG_O_ and ratBG_O_ highlighted the importance of the BL of ratBG_O_ not only for efficient binding to TGF-βs, and thus indirect potentiation of binding of the signaling receptors by increasing the local concentration, but also for direct potentiation of TGFBR2 binding. This dual mechanism allows for more equal binding of all TGF-β isoforms to the signaling receptors.

**Figure 5.**
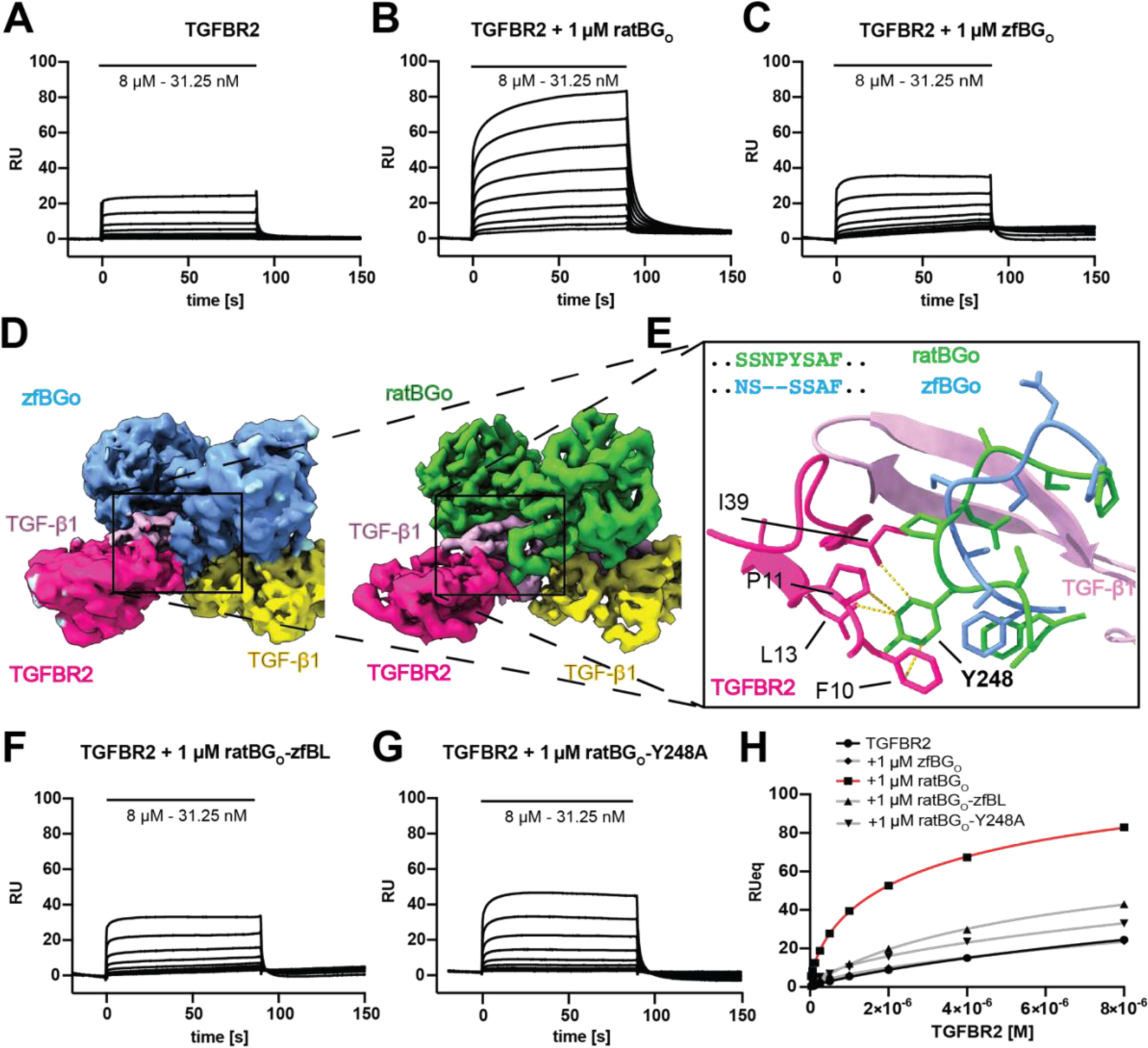
ratBGo directly contacts and potentiates binding of TGFBR2. (A–C) SPR sensorgrams showing binding of TGFBR2 to immobilized TGF-β2. The same experiment was repeated in the presence of 1 µM ratBG_O_ (B) or 1 µM zfBG_O_ (C). (D) Zoom into TGF-β binding loop regions of CryoEM strucutres of zfBGo and ratBG_O_ in complex with TGF-β1 dimer and two TGFBR2 ECDs. (E) A blow-up of the boxed area is demonstrating how ratBGo, but not zfBGo, can form additional contact with TGFBR2 through Tyr^248^. (F, G) SPR sensorgrams showing binding of TGFBR2 to the immobilized TGF-β2 in the presence of 1 µM ratBG_O_-zfBL (F) and 1µM ratBG_O_-Y248A (G). (H) Plot of the normalized equilibrium responses of TGFBR2 binding to TGF-β2 in the presence of different variants of ratBG_O_.

### InhA competes with TGF-**β** through binding to distinct but overlapping regions of BG_ZP–C_

InhA and InhB play crucial roles in endocrine signaling pathways that regulate fertility^19^. InhA and InhB are heterodimers which share a common α-subunit but differ in their β-subunit composition. InhA contains a βA subunit, which is also shared with ActA, while InhB contains a βB subunit, which is shared with activin B (ActB). InhA binds the activin type II receptors, ACVR2 or ACVR2B, through its βA subunit. In cells, this interaction is further stabilized by the binding of BG_ZP-C_ via the α-subunit, effectively sequestering ACVR2 or ACVR2B from forming a signaling complex with ActA and the activin type I receptor, ACVR1B. Despite sharing the same α subunit with InhA, InhB has only a modest binding affinity for the BG_ZP-C_ domain^28^. Recent studies have shown that InhB primarily exerts its effects in gonadotrope cells through a TGFBR3-like, TGFBRL, diverging from the pathway utilized by InhA^28^.

These observations strongly indicate that both the α and β subunits of InhA mediate binding to BG_ZP-C_. We therefore attempted to fit InhA into the TGF-β binding site such that the α subunit of InhA aligns similarly to one monomer of TGF-β2. In this arrangement, the β subunit would be able to contribute to the binding process in a manner comparable to how the second monomer of TGF-β2 interacts, specifically through the heel helix and FG-helix of BG_ZP-C_. To test our hypothesis, we employed Nuclear Magnetic Resonance (NMR) spectroscopy, focusing on the BG_ZP-C_ domain with^13^ C-methyl-Ile labelling. This technique allows detection of distinct ’fingerprints’ of the BG_ZP-C_ domain both unbound and bound to the ligand, as depicted in Fig. 6A-B. Our data show that both the TGF-β2 dimer and mmTGF-β2 generate highly similar patterns of chemical shift changes, including the transition of Ile^161^ from a random coil to a non-random coil chemical shift, consistent with the disordered loop to helix transition observed in the X-ray structure.

**Figure 6.**
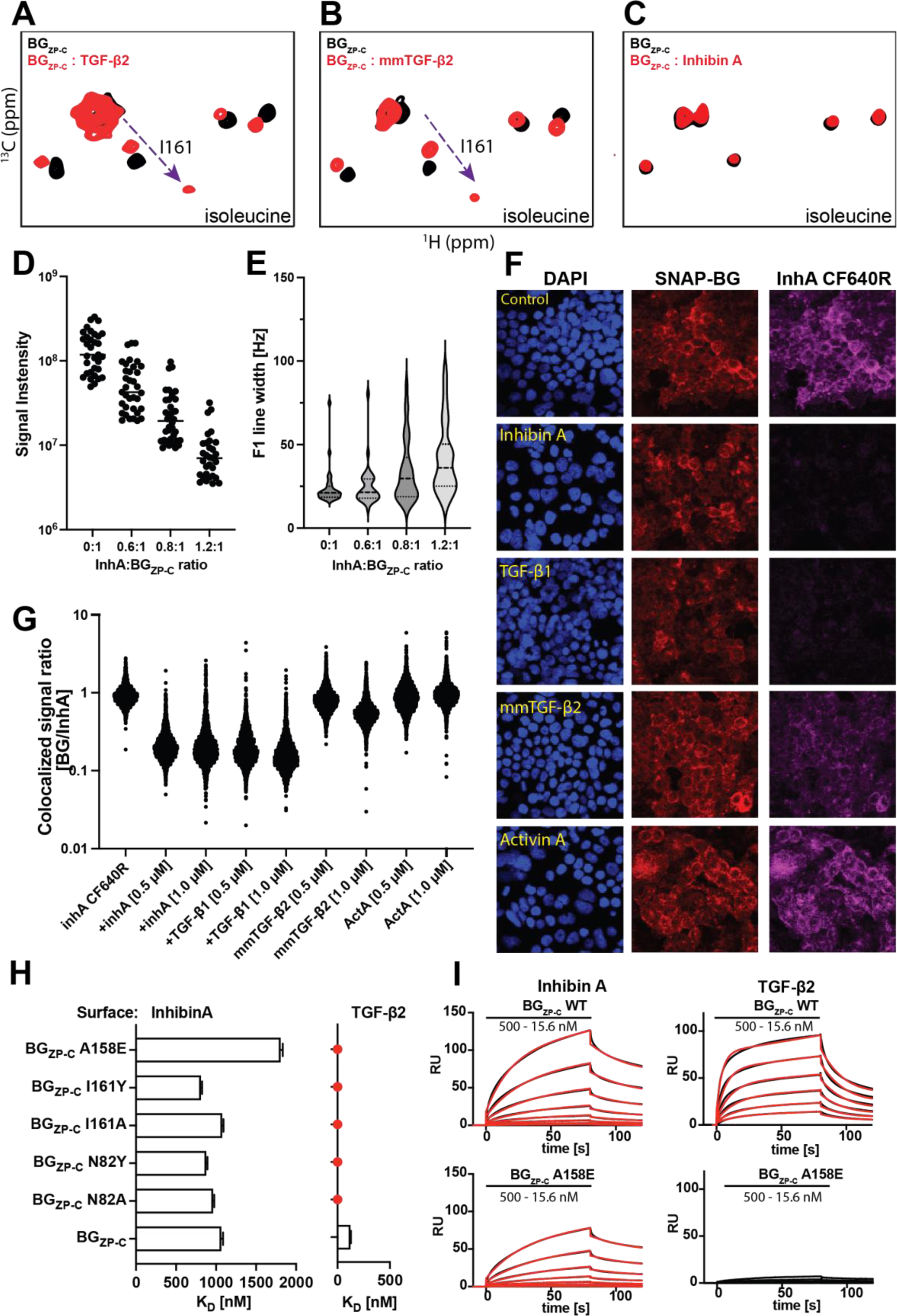
InhA and TGF-β interact with partially overlapping, but distinct regions of BG_ZP-C_. (A-C) Isoleucine δ1 methyl region of the ^1^H-^13^C correlation NMR spectra of BG_ZP-C_ (black). Upon TGF-β (A) or mmTGF-β2 (B) binding (red), the chemical shift of Ile^161^ δ1 methyl is perturbed, with the signal shifting from the random coil to the structured region, consistent with disordered loop to helix transition. No shifts were observed upon binding of InhA (C). (D) Signal intensity and (E) line width change of Ile δ1 signals upon InhA titration into methyl labeled BG_ZP-C_ sample. (F) Confocal microscopy images of competition assay - HEK293T cells stably transfected with a SNAP-tagged BG construct (SNAP-BG) stained with DAPI (left panels), incubated with a SNAP-tag ligand conjugated to the CF567 fluorescent dye (middle panels), along with InhA tagged with CF640R fluorescent dye (right panels) alone (row 1) or with the addition of the indicated unlabeled ligands (rows 2-5). Bound InhA could be outcompeted by addition of excess unlabeled InhA, TGF-β2 or mmTGF- β2. (G) Quantification of the fluorescent colocalized SNAP-BG/InhA signals from microscopy data. (H) Chart summarizing SPR binding experiments using selected panels of BG_ZP-C_ mutants binding to immobilized TGF-β2 or InhA. (I) Selected SPR binding sensorgrams of BG_ZP-C_ mutants binding to immobilised TGF- β2 or InhA on the chip surface.

Upon titration of ^13^C-methyl-Ile BG_ZP-C_ with InhA, we observed a concentration-dependent weakening of signal intensities and peak broadening, which is indicative of binding (Fig. 6C-E). However, the exact cause of the peak broadening remained unclear, making it challenging to draw definitive conclusions about the position of the binding interface. Therefore, we conducted a cell-based competition experiment in which InhA and TGF-β directly compete for binding to the BG_ZP-C_ domain (Fig. 6F, G). In this experiment, we labelled InhA with a fluorescent marker and observed clear co- localization with SNAP-BG, which was stably expressed in our HEK293 cell line (Fig. 6F, top). This co-localization was disrupted by the addition of an excess of unlabeled InhA or TGF-β1, indicating that InhA forms a complex with SNAP-BG and that InhA and TGF-β compete for binding to BG_ZP-C_ (Fig. 6F, rows 2-3). mmTGF-β2 also disrupted the co-localization, though consistent with its reduced binding affinity for BG_ZP-C_, it competed with reduced potency (Fig. 6F, row 4) and required a higher ratio to obtain similar results (Extended Data Fig. 6C). ActA did not diminish the co- localization of InhA with BG_ZP-C_, consistent with its inability to bind BG_ZP-C_ (Fig. 6F, row 5).

In addition, we also investigated how mutation of specific residues on BG_ZP-C_ effect binding to InhA relative to TGF-β (Fig. 6H). Interestingly, mutations such as Asn^82^ or Ile^161^ to Ala, and even more severe alterations, such as Asn^82^ or Ile^161^ to Tyr, that disrupted binding to TGF-β2 did not significantly affect binding to InhA. The only mutation that caused a modest reduction in binding was Ala^158^ to Glu (Fig. 6H and Extended Data Fig. 6A-B). Collectively, these results indicate that InhA binds to BG_ZP-C_ through an interface that partially overlaps with the one determined for TGF-β. However, the exact position or involvement of structural features on BG_ZP-C_ that enable binding likely differ.

## Discussion

Here, we present the structure of full-length BG bound to TGF-β, offering new insights into its specificity of ligand binding and mechanism of signal potentiation. We combined X-ray crystallography, Cryo-EM, and NMR spectroscopy that allowed us to study the binding interfaces in fine detail, in the context of the full-length co-receptor and in the presence of the signaling receptors. The determined structures show how the BG_O_ and BG_ZP-C_ domains bind independently to TGF-β dimers, allowing for efficient recruitment of TGF-β to the cell surface and how the signaling receptors take over the ligand and thus signal (Fig. 7A). Using our cell-based fluorescent assay and advanced modeling with AF2M, we also showed the remarkable BG specificity towards specific members of TGF-β ligand family (Fig. 7B) and using both site-directed mutagenesis and modeling we demonstrate how this specificity is achieved. Lastly, leveraging structural, biophysical, and cell-based assays we also provide insights into how BG engages InhA, which is distinct from the TGF-β isoforms, however InhA competes with them for binding (Fig. 7C).

**Figure 7.**
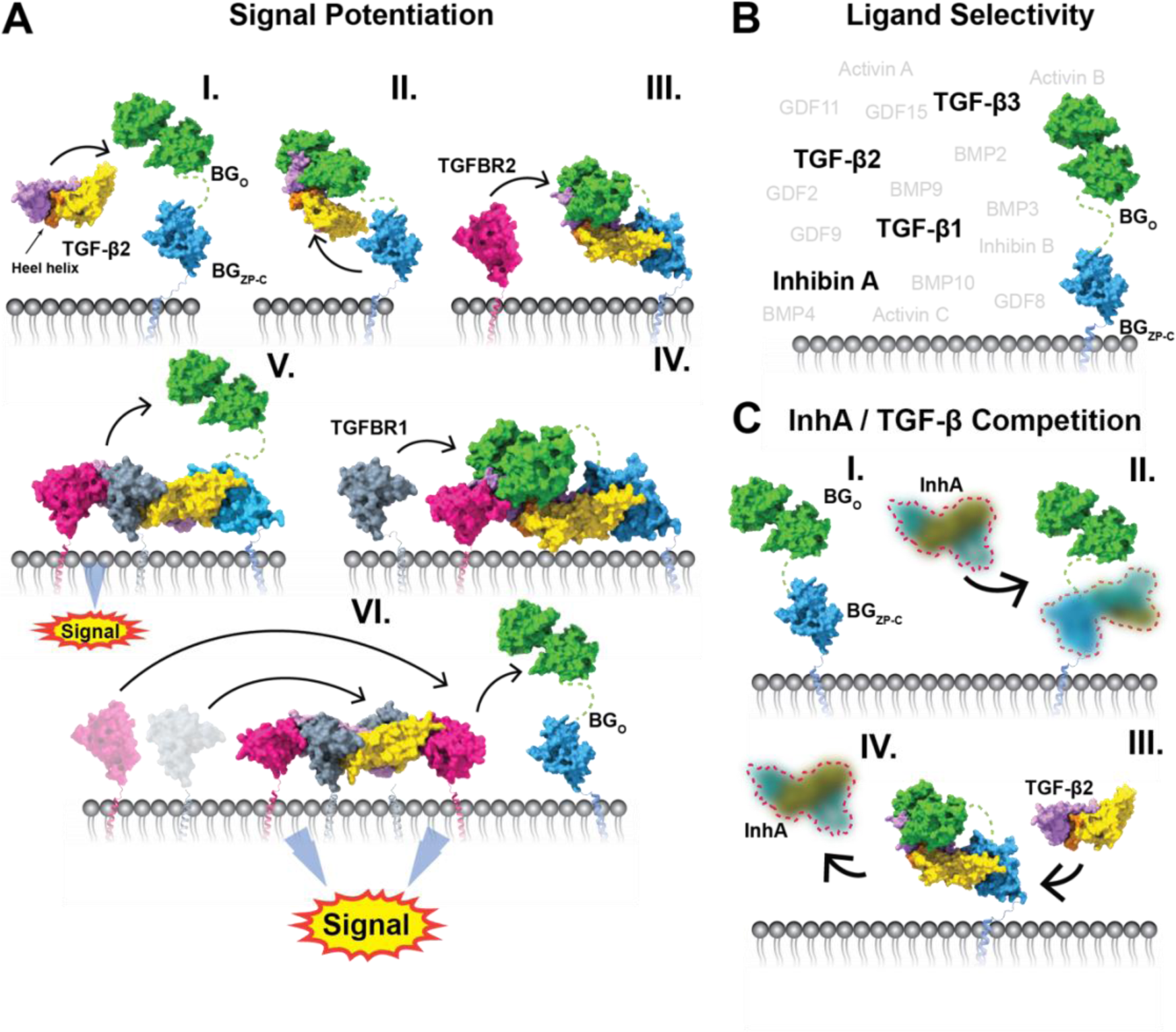
Structural illustration of BG ligand selectivity and TGF-β2 signaling enhancement. (A) Mechanism of TGF-β2 signal potentiation by BG. In the first step TGF-β2 binds to BG_O_ (I), then the BG_ZP-C_ domain binds (II). The presence of BG_O_ increases the affinity towards TGFBR2 (III) that binds to the available site adjacent to the BG_O_ domain. Next, TGFBR1 is recruited by binding to the shared TGF-β2:TGFBR2 interface (IV), which displaces BG_O_ (V). This complex can already signal, however BG_ZP-C_ can be further displaced and the heterotetrameric signalling complex, with full signaling potential,can be formed (VI) (B) Schematic showing BG ligand selectivity towards TGF-β isoforms as well as InhA among other TGF-β family members. (C) InhA competes with TGF-β2 for binding to BG_ZP-C_. BG_ZP-C_ binds to InhA (blurred picture denotes lack of determined molecular structure) and forms a complex (I, II). TGF-βs can compete off InhA from its binding position (III, IV).

### The mechanism by which BG potentiates TGF-β signaling

The high-resolution structures of the zfBG_O_ and BG_ZP-C_ domains bound to TGF-β, together with the low-resolution structure of the TGF-β1:BG complex, show that BG_O_ binds TGF-β such that the adjacent TGFBR2 binding site is still accessible for binding. In contrast, the BG_ZP-C_ domain blocks the binding site of both TGFBR1 and TGFBR2. Thus, once BG binds and captures TGF-β ligands on the cell surface, this likely initiates the recruitment of one TGFBR2 at the one receptor binding site that remains accessible (Fig. 7A). The structure of the TGF-β1:BG complex further shows that BG_ZP-N_ neither participates in binding nor is rigidly tethered to BG_ZP-C_. This explains indirect observations that the BG_ZP-N_ domain is not required for TGF-β potentiation^4^

The structural and biophysical binding data demonstrate that the binding loop (BL) of ratBG_O_ is not only the main structural element required for binding to the TGF-βs, but also potentiates the binding of TGFBR2 through direct co-receptor–receptor contact. The potentiation is mediated by expanding the binding interface and increasing the affinity, not through allostery. The two amino acid deletion occurring in *Danio rerio* BG completely impairs the direct, yet not indirect, potentiation of TGFBR2 binding and is also present in most other boney fish families. In contrast, the extended version is present not only in amphibians, reptiles, birds, and mammals, but also in cartilaginous fish and primitive bone fish from the *Polyodontidae*, *Acipenseridae*, and *Polypteridae* families. This pattern suggests a divergent evolution of the BG protein in teleosts, rendering it a possibly less transferable model for studying TGF-β signaling by BG in other vertebrates, as previously noted^53^.

The transition from the TGF-β:BG:TGFBR2 ternary complex to the signaling complex has been proposed to occur by a handoff mechanism, wherein the one bound TGFBR2 potentiates the recruitment of TGFBR1 through direct receptor–receptor contact, and in the process displaces the orphan domain, leaving the second monomer of TGF-β bound by only the BG_ZP-C_ domain (Fig. 7A)^4, 8^. Inspection of the TGF- β:BG_O_:(TGFBR2)_2_ and the TGF-β3WD:zfBG_O_:TGFBR2:TGFBR1 structures shows that the BL of BG_O_ partially occupies the TGFBR1 interface formed by the heel helix of the adjacent TGF-β monomer and TGFBR2, which would be expected to block binding of TGFBR1. However, the binding affinity of TGFBR1 to the TGF-β:TGFBR2 binary complex (K_D_ ca. 30 nM)^41^ is significantly higher than BG_O_ binding to TGF-β (K_D_ ca. 200 nM)^8^, enabling TGFBR1 to outcompete and displace BG_O_. It is notable that the heel helix of the second monomer of TGF-β also takes part in binding of BG_O_, albeit through a limited interface formed mostly by two residues, Asp^55^, Thr^57^ (Fig. 4A III, D), located on the solvent-exposed surface of the heel helix. Mutations in this region of TGF-β2 almost completely abrogate BG_O_ binding (Fig. 3E and Extended Data Fig. 3A) and are likely responsible for the diminished binding of mmTGF-β2 (ref 26). This suggests that relatively small changes close to the heel helix may have a pronounced effect on BG_O_ binding and thus partial docking of TGFBR1 may facilitate the rapid release of BG_O_ that drives the handoff (Fig 7A IV-V).

However, it is unclear how TGFBR2 or TGFBR1 can outcompete BG_ZP-C_ from its binding position. The structure of the mmTGF-β2:BG_ZP-C_ complex shows that BG_ZP-C_ not only binds and engages the two critical basic residues on the tips of fingertips where TGFBR2 binds, but also occupies the underside of the fingers where TGFBR1 binds. In modelling studies it was shown that in spite of the lower affinity of TGFBR2 and TGFBR1 for binding TGF-β2 compared to BG_ZP-C_, together these can nonetheless displace BG_ZP-C_ to form the TGF-β2(TGFBR2)_2_(TGFBR1)_2_ signaling complex, due to transient disassociation of BG_ZP-C_ and sequential binding of TGFBR2 then TGFBR1 or TGFBR1 then TGFBR2 (ref 38). There are two mechanisms that could further promote this manner of assembly. The first is internalization of the TGF- β2:BG:TGFBR1:TGFBR2 complex into endosomes, which may occur based on a previous report that the disordered but highly conserved cytoplasmic tail of BG is bound by β-arrestin and facilities internalization^54^. Then, assembly might follow a mechanism demonstrated for the repulsive guidance mediators (RGMs), co-receptors for some BMPs, whereby internalization in endosomes triggers the disassociation of the co-receptor due to a lowering of the pH, followed by binding of the signaling receptors. The second possible mechanism is that proteolytic cleavage might occur within the BG_ZP-C_ FG-loop^11^ thereby weakening binding and enabling recruitment of a second pair of signaling receptors (Fig. 7A VI).

It is also possible that the BG is not fully displaced and the ligand signals as a quaternary complex with BG_ZP-C_ still bound. This is suggested by the finding that TGF- βs bound to a single TGFBR2:TGFBR1 heterodimer can signal. However, in the aforementioned modeling studies^38^, this manner of binding could not recapitulate the biphasic effect on TGF-β2 signaling, wherein low density BG potentiates signaling, while at higher density BG has the opposite effect. Even though the biphasic behavior is well supported by experimental data, there is no direct experimental evidence showing a transition point where BG would turn from potentiation to inhibition as a function of BG density. Therefore, the second phase where BG diminishes TGF-β2 signaling could be attributed to domain shedding^55^. This explanation is further suggested by the detection of BG:TGF-β2:TGFBR2 complexes in crosslinking experiments^4^, suggesting the non-transient nature of this complex.

### BG ligand binding specificity

BG demonstrated remarkable selectivity towards the three TGF-β isoforms and InhA as assessed by our competition assays. Modeling with AF2M allowed us to broaden our investigation to include interactions between BG and all ligands of the TGF-β family. The experimental data and predictions from AF2M identified only the three TGF-β isoforms as binding partners of BG_O_. Inspection of the binding interface, together with the multiple sequence alignment (MSA), highlighted residues likely responsible for ligand selectivity. However, we were not able to evaluate their significance using mutagenesis as substitution of these amino acids disrupted the protein structure. In spite of this, their importance was confirmed by AF2M, indicating that these residues are likely important determinants of the specificity.

On the BG_ZP-C_ domain, the key residues suggested by the mmTGF-β:BG_ZP-C_ structure, such as Asn^82^ and Ile^161^, were shown to be essential for binding, though most of the hydrophilic interactions occur between the side chain of one of the partners and the backbone of the other. The backbone atoms of closely related isoforms are expected to have a more consistent arrangement than the sidechains, which may enable the co-receptor to form stable interactions with more than one partner that share the same geometry but differ in sequence. The previously reported binding site residues that differ from those demonstrated by our structure^30, 34^ likely arose due to insufficiently validated mutants, as some of the mutations likely impacted folding and/or trafficking to the membrane, rather than directly impacting ligand binding.

BG_ZP-C_ is known to bind all TGF-β isoforms, but also InhA^18, 26, 28^. InhA binds to BG_ZP-C_ through an interface that at least partially overlaps with the TGF-β interface, yet binding of InhA to BG_ZP-C_ is insensitive to most of the mutations that abrogate binding to TGF-β. This suggests that BG_ZP-C_ engages InhA differently than it does with TGF-β, though the precise manner of binding is unclear, as models of the InhA:BG_ZP-C_ complex generated using AF2M were variable and could not simultaneously account for both the competition and mutagenesis data. It is notable that for AF2M predictions of BG_ZP-C_ complexes, the score gap between binders and non-binders is significantly narrower compared to the predictions for BG_O,_ lowering the confidence of the predictions. Moreover, no structure of InhA is available, thus modeling is expected to be more challenging, and obtaining a structure of the InhA:BG_ZP-C_ complex will be essential. Our work does not rule out the possibility that BG can engage some other ligands of the TGF-β family, as suggested by some published data^44, 45, 56^. However, if such complexes form, these are likely significantly weaker than the complexes formed with the TGF-β isoforms or involve interfaces different from those described here.

### Structural insights into TGF-β pathway inhibition

The TGF-β pathway has been pharmacologically targeted using several strategies, including small molecule receptor kinase inhibitors, peptides, antisense oligonucleotides, neutralizing antibodies and ligand traps based on fusing of the extracellular domains of the TGF-β receptors. It is worth noting that TGFBR2-based traps, such as AVID200 (ref ^57^) or Bintrafusp Alfa^58^ or neutralizing antibodies, such as GC1008 or GC2008 (ref ^59^), would be expected to compete for binding with membrane bound BG and demonstrate attenuated efficacy in environments highly enriched with BG. Indeed, during our structural studies we attempted to utilize the neutralizing antibody GC2008 to increase orientational sampling for Cryo-EM. However, it was not possible to obtain a complex with both BG and GC2008 bound to TGF-β, as the components competed with one another.

One interesting strategy of inhibiting TGF-β is the use of a short, receptor-derived peptide that would block the interaction of the ligand with the receptor. One such example is the 14-mer peptide P144, whose sequence is derived from the BG_ZP-C_ domain of human BG. It was shown that P144 blocks TGF-β1 biological activity in different *in vitro* and *in vivo* models^35, 60–62^ and it currently is the drug candidate Disitertide®, which is being tested for pathological skin fibrosis, such as scleroderma. Indeed, our mmTGF-β2:BG_ZP-C_ structure demonstrates that the sequence is precisely derived from the loop region that forms the FG-helix upon binding to concave surface of the fingers of the growth factor^60^. We anticipate that our structural data will advance the knowledge-based design of next-generation inhibitors, enhancing both their potency and biochemical properties.

## Acknowledgments

We would like to thank Simone Kunzleman and Chloe Roustan from the Structural Biology STP at the Crick for assistance with SPR experiments and HEK293T protein expression and Hema Nagaraj from the Chemical Biology STP for providing fluorescently labelled SNAP-ligand. We would also like to thank Mateo Coppola who helped with the production of the BG_O_ mutants, Jue Wang from DeepMind for useful discussion, Scott Wilcockson for his helpful comments and corrections, James Conway from the CryoEM laboratory at the University of Pittsburgh and Daniel Clare from UK national electron Bio-Imaging Centre for support in microscope alignment and data collection. This research was supported by the NIH through RO1 grants (GM58670 and CA233622) awarded to A.H. and the CIHR through a grant (PJT- 191766) awarded to D.J.B. This work was also supported by the Francis Crick Institute, which receives its core funding from Cancer Research UK (CC2021, CC2058), the UK Medical Research Council (CC2021, CC2058), and the Wellcome Trust (CC2021, CC2058) to C.S.H. and P.C. This project additionally received funding from the European Union’s Horizon 2020 research and innovation programme under the Marie Skłodowska-Curie Grant Agreement No. 893196 to L.W. This work is based upon research conducted in the Structural Biology Core, a part of the Institutional Research Cores at the University of Texas Health Science Center at San Antonio supported by the Office of the Vice President for Research, Greehey Children’s Cancer Research Institute, and the Mays Cancer Center Drug Discovery and Structural Biology Shared Resource (NIH P30 CA054174). The Rigaku HyPix-6000HE Detector, Universal Goniometer and VariMax-VHF Optic instrumentation are funded by NIH-ORIP SIG Grant S10 OD030374. X-ray data were collected at Northeastern Collaborative Access Team (NECAT) 24-ID beamline at the Advanced Photon Source, Argonne National Laboratory supported by the NIH (GM124165, RR029205, and OD021527) and the DOE (DE-AC02-06CH11357). Cryo-EM data was collected at the Cryo-EM laboratory at the University of Pittsburgh supported by the NIH (S10 OD025009) and at the UK national electron Bio-Imaging Centre (eBIC) under proposal BI37190.

## METHODS

### Cloning

All constructs used to express proteins for X-ray Crystallography and CryoEM were already available from previous studies (Supplementary Table 4). For SPR binding studies a set of mutants were prepared using site-directed mutagenesis of pCDNA3.1+-ALB-BG_ZP-C_-(N82A, N82Y, K86A, D88A, E130A, L143A, S155A, A158A, A158E, I161A, I161Y, W162A, M164A, Q166A); pCDNA3.1+-zfBG_O_-(F49A, L208A, L210A, L213A, F250A, D253A, K316A, K319A) or by cloning respective GenBlocks into pCDNA3.1+ vector (IDT DNA) pCDNA3.1-ratBG_O_-zfBL, pCDNA3.1-ratBG_O_- Y248A, pCDNA3.1-ratBG_O_-BL, pCDNA3.1+-proTGF-β2, pCDNA3.1+-proTGF-β2- (E99K, L101Y, S102Q, N103D), pCDNA3.1+-proTGF-β2-(E99K, S102Q, N103D), pCDNA3.1+-proTGF-β2-(E99K, S102Q), pCDNA3.1+-proTGF-β2-(S102Q, N103D), or into pET32a vector pET32a-TGF-β2-(Q57A, K60A, L64A), pET32a-TGF-β2-(Y50A, L51A) using In-Fusion seamless cloning (Takara Bio). SNAP-BG was constructed using overlapping PCR, joining SNAP domain with c-terminal linker “GGGGSGGGGSGGG” to N-terminus of rat BG (residues 30..853) and cloning entire construct to pCDNA3.1+ vector. The CAGA_12_-luciferase reporter^47^ and the BG_ZP-C_ construct^20^ used in luciferase assays were described previously. The mutant constructs were constructed using the QuikChange protocol with the following primers: N82A_Forward-TGATTACACCATCATCGAGGCCATCTGTCCGAAAGACGAC, N82A_Reversed-GTCGTCTTTCGGACAGATGGCCTCGATGATGGTGTAATCA, I161A_Forward-CTCTCGATGCCACCATGGCCTGGACCATGATGCAGA, I161A_Reversed-TCTGCATCATGGTCCAGGCCATGGTGGCATCGAGAG.

### Expression and purification of TGF-β growth factors

The growth factor domains of human TGF-β2, TGF-β3, TGF-β3WD and mmTGF-β2 were produced as described before^39^. Briefly, proteins were expressed in *E. coli* BL21(DE3) in the form of insoluble inclusion bodies and after washing, the material was reconstituted in 8 M urea, reduced using DTT and refolded by dilution into non-denaturing buffer. The natively folded growth factors were isolated by high-resolution cation–exchange chromatography using MonoS column. Intact masses of the purified proteins were measured by LC electrospray ionization TOF MS (LC-ESI-TOF-MS, Bruker Micro TOF, Billerica, MA) and folding was assessed by 2D NMR. Pro-TGF-β1 was produced in HEK293 suspension culture and mature growth factor was purified from pro-domain by high-resolution cation–exchange chromatography on MonoS column (GE Healthcare).

### Expression and purification of mammalian-expressed BG_ZP-C_ and BG_O_

Rat betaglycan extracellular domain BG_ZP-C_ and the rat or zebrafish orphan domains, (ratBGO, zfBG_O_, respectively) were expressed as secreted proteins in suspension-cultured expi293F cells and expi293^TM^ expression medium, as described previously^8^. To gain better yield, the BG_ZP-C_ domain was expressed fused to albumin, which was removed by proteolytic cleavage with thrombin and separated by size exclusion chromatography.

### Expression and purification of bacterially expressed BG_ZP-C_ domain

An isotopically labeled sample of BG_ZP-C_ for NMR studies was produced in *E. coli* BL21 (DE3) in form of insoluble inclusion bodies and were refolded and purified as previously described in detail^63^.

### Expression and purification of Inhibin A

Inhibin A heterodimer was expressed as a secreted protein in suspension cultured expi293F cells and expi293^TM^ expression medium. Heterodimer was purified using in tandem -His and -Strep tags present on separate chains and further polished using reverse phase chromatography. Protein activity was determined by cell activity assay as previously reported^28^.

### Protein folding assessment by NMR

Proper folding of selected mutant proteins was assessed by recording NMR natural abundance methyl-TROSY ^1^H-^13^C correlation spectra (in case of BG_ZP-C_) or 1D proton spectra (in case of zfBG_O_) and by comparing the spectral fingerprints to the wild type proteins. NMR spectra were recorded with protein samples concentrated to at least 150 µM in 25 mM phosphate buffer pH 6.0 and using a band-selective optimized-flip-angle short-transient experiment^64^ (SOFAST-HMQC) or 1D proton Bruker experiment with Watergate water suppression sequence. All measurements were recorded at 37°C at 600 or 700 MHz Bruker spectrometer equipped with a Bruker TCI triple-resonance cryogenically cooled probes running Topspin v2.0 or v3.5. Data was processed with NMRPipe^65^ and analyzed with CcpNmr Analysis software package^66^.

### NMR Methyl spectra of BG_ZP-C_: Growth Factor binding

Samples of ^2^H, ^13^C-methyl Isoleucine, Leucine, Valine (ILV) labelled BG_ZP-C_ were prepared in 25 mM ^2^H-glycine buffer at pH 11.0 or 25 mM phosphate buffer at pH 6.0 in either straight 3 mm tubes (Wilmad, Vineland, NJ) or 5 mm susceptibility-matched microcells (Shigemi, Sigma-Aldrich). Separate samples, each with the labelled protein being 15 – 25 μM, were used for the titration. Spectra were recorded at 37°C using a band-selective optimized-flip-angle short-transient experiment (SOFAST-HMQC)^64^. All measurements were recorded at 37°C at 600 MHz Bruker spectrometer equipped with a Bruker TCI triple-resonance cryogenically cooled probes running Topspin v3.5. Data was processed with NMRPipe^65^ and analyzed with CcpNmr Analysis software package^66^.

### TGF-β2 biotinylation for SPR

TGF-β2 and its variants were dialyzed into 10 mM acetic acid and 125 µg aliquots were transferred into Eppendorf tubes and lyophilized overnight. Samples were reconstituted in 4 mM HCl. In order to remove traces of acetic acid, lyophilization was repeated and samples were resuspended in 50 µl of 4 mM HCl. 300 µL of 25 mM MES, pH 6.0. + 40% DMSO buffer was added, followed by 0.0115 mg of ethyl-3-(3- dimethylaminopropyl)carbodiimide hydrochloride (EDC, Pierce) (60 nmol), 0.026 mg of sulfo-N-hydroxysulfosuccinimide (12 nmol, Pierce), and 0.20 mg (+)-biotinyl-3,6,9- trioxaundecanediamine (EZ-Link Amine-PEG3-Biotin, 480 nmol, Pierce). The reaction was allowed to proceed for 2h at room temperature and then quenched by the addition of 1 mL 100 mM acetic acid. The growth factor was repurified by loading it onto a MonoS high resolution cation exchange column equilibrated in 25 mM sodium acetate, 30% isopropanol pH 4.5 and eluting with salt using gradient from 0 – 0.7 M over 10 column volumes. Modification of the ligand was confirmed by LC-ESI-TOF MS.

### SPR measurements of BG_ZP-C_ mutants

Binding studies of BG_ZP-C_ or zfBG_O_ mutants were done using streptavidin-coated sensor chips for capture of biotinylated ligands. Surface of a CM-5 sensor chip (GE Healthcare) was activated with EDC and N-hydroxysulfosuccinimide (NHS) (Pierce) followed by injection of streptavidin (Pierce) diluted into sodium acetate at pH 4.5 until the surface density reached 6000 - 8000 RUs. Biotinylated ligands were captured onto the streptavidin surface to a surface density between 50–300 RU. Equilibrium binding assays were performed by injecting the analytes in 10 mM HEPES, 150 mM NaCl, 0.1% surfactant P20 (Pierce) at pH 7.4 (HBS-EP buffer) at a rate of 10 μL min^−1^ for 720 s, followed by a dissociation period of 600 s. Regeneration of the surface was achieved by a 10 s injection at 100 μL min^−1^ of 4 M guanidine hydrochloride in 4 mM HCl solution. Baseline correction was performed by double referencing. The data was analyzed by fitting the results to a 1:1 kinetic model using the SPR analysis software Scrubber (BioLogic Software). Details of binding parameters are listed in Supplementary Table 3.

### SPR measurements of zfBG_O_ mutants

SPR experiments were performed with a BIAcore X100 system (Cytiva). Neutravidin was coupled to the surface of a CM5 chip (Cytiva) by EDC-NHS activation of the chip, followed by injection of neutravidin (Thermo) over the surface in sodium acetate, pH 4.5 until the RU increased by 10000-15000 RU. Minimally biotinylated TGF-β2 was prepared by NHS/EDC activation followed by addition of EZ-Link Amine-PEG3-Biotin (Thermo). After removal of reagents, biotin-TGF-β2 was captured onto the chip surface at a maximum density of 50 RU. All experiments were performed in 10 mM CHES, 150 mM NaCl, 3 mM EDTA, 0.01% P20 surfactant, pH 8.6 at an injection rate of 100 μl min^-1^. The surface was regenerated in between each injection with a 10 second injection of 0.2 M guanidine hydrochloride. The experimental sensograms were obtained with double referencing with a control surface coated similarly with neutravidin but lacking the captured TGF-β2 and 8 blank buffer injections at the beginning of the run before injection of the samples. The data was analyzed by fitting the results to a 1:1 kinetic model using the SPR analysis software Scrubber (BioLogic Software). Details of binding parameters are listed in Supplementary Table 3.

### SPR measurements of BG_O_ potentiation of TGFBR2 binding

SPR binding studies were performed on BIAcore T200 (Cytiva) and analyzed with the Biacore T200 analysis Software v3.0 at the Francis Crick Institute. TGF-β2 was coupled to the surface of a CM4 chip (Cytiva) by EDC-NHS activation of the chip, followed by injection of 1 µM TGF-β2 over the surface in sodium acetate, pH 4.5 until the RU increased by maximum 400 RU. All experiments were performed in 10 mM CHES, 150 mM NaCl, 3 mM EDTA, 0.01% P20 surfactant, pH 8.0. The A-B-A injection scheme was applied at an injection rate of 30 μL min^-1^ in sequence 60-90-60 s. The surface was regenerated in between each injection with a 10 second injection of 0.2 M guanidine hydrochloride pH 2.5. The experimental sensorgrams were obtained with double referencing with a control surface and 4 blank buffer injections. The data was analyzed by fitting the results to a 1:1 kinetic model using the SPR analysis evaluation tool (Cytiva).

### SPR measurements of additional mutants of BG_ZP-C_ (N82Y, I161Y, A158E) and TGF-β2 mutants (Q57A, K60A, L64A and Y50A, L51A)

SPR binding studies were performed on BIAcore T200 (Cytiva) and analyzed with the BIAcore T200 analysis Software v3.0 at the Francis Crick Institute. TGF-β2 or TGF- β2 mutants (Q57A, K60A, L64A and Y50A, L51A) were coupled to the surface of a CM4 chip (Cytiva) by EDC-NHS activation of the chip, followed by injection of 1 µM ligand over the surface in sodium acetate, pH 4.5 until the RU increased by maximum 400 RU. All experiments were performed in 10 mM CHES, 150 mM NaCl, 3 mM EDTA, 0.01% P20 surfactant, pH 8.0. injection rate of 100 μL min^-1^. The surface was regenerated in between each injection with a 10 s injection of 0.2 M guanidine hydrochloride. The experimental sensorgrams were obtained with double referencing with a control surface and 4 blank buffer injections. The data was analyzed by fitting the results to a 1:1 kinetic model with drift using the SPR analysis evaluation tool (Cytiva). Details of binding parameters are listed in Supplementary Table 3.

### Promoter-reporter assays

L6E9 cells were seeded at a density of 20,000 cells/well in a 48 well plate. The following day, cells were transfected with the CAGA_12_-luciferase reporter (200 ng/well) and either empty vector or a BG_ZP-C_ construct (100 ng/well) using Lipofectamine 3000, following the manufacturer’s protocol. Twenty-four hours after transfection, cells were serum starved for an additional 24 hours. After starvation, cells were lysed in 50 µL/well passive lysis buffer (25 mM Tris-phosphate [pH 7.8], 10% [v/v] glycerol, 1% [v/v] Triton X-100, 1 mg/mL bovine serum albumin, 2 mM EDTA) for 10 minutes at room temperature with agitation. Twenty microliters of cell lysis supernatant were combined with 100 µL of assay buffer (15 mM potassium phosphate [pH 7.8], 25 mM glycylglycine, 15 mM MgSO4, 4 mM EDTA, 2 mM adenosine triphosphate, 1 mM dithiothreitol, 0.04 mM D-luciferin), and luciferase activity was measured on an Orion II microplate luminometer (Berthold Detection Systems, Oak Ridge, TN, USA).

### Crystallization, structure determination and refinement

Automated screening for crystallization was carried out using the sitting drop vapor-diffusion method with an Art Robbins Instruments Phoenix system in the Structural Biology Core at The University of Texas Health Science Center at San Antonio. Crystals of mmTGF-β2:BG_ZP-C_ were grown at 22°C in 13% PEG 4000, 0.1 M sodium citrate pH 5.6 and 10% ethylene glycol mixed in a 1:1 ratio for a total drop volume of 0.4 µL. Likewise, crystals of TGF-β3WD:BG_O_:TGFBR1:TGFBR2 were grown at 22°C in 30% PEG 4000, 0.1 M tris pH 8.5 and 0.2 M lithium sulfate. Diffraction data were collected at the Advanced Photon Source, Argonne, IL, NE-CAT beamlines for crystals flash-cooled in liquid nitrogen after wicking off excess solution from the crystals harvested in nylon cryo-loops. Data were processed using AUTOPROC^67^. The structure of mmTGF-β2:BG_ZP-C_ was determined by the molecular replacement method implemented in PHASER ^68^ using coordinates from PDB entries 3QW9^34^ and 5TX6^39^ as search models. The structure of TGF-β3WD:BG_O_:TGFBR1:TGFBR2 was determined using coordinates from PDB entries 2PJY^42^ and 6MZN^22^ as search models. A successful molecular replacement phase solution was achieved for this complex when searching with separate OD1 and OD2 subdomains for the BG_O_ component. All coordinates were refined using PHENIX^69^ with simulated annealing and TLS refinement, and alternated with manual rebuilding steps using COOT^70^. The models were verified using composite omit map analysis^71^. Data collection and refinement statistics are shown in Supplementary Table 1.

### CryoEM analysis of TGF-β1:zfBGo:(TGFBR2)_2_ complex and model refinement

Aliquots of 3 μL of each complex were applied to glow discharged UltrAuFoil r1.2/1.3 grids. The grids were blotted for 3 seconds at 100% humidity with force “2” and plunge frozen into liquid ethane using a Vitrobot Mark IV (Thermo Fisher). 41335 movies were recorded on a 300 keV Titan Krios microscope (Thermo Fisher) equipped with a FalconIV camera and a Selectris energy filter using 0.72 Å/pix. The total dose was set to 50 electrons per Å^2^. Movies were corrected using patch motion correction in CryoSPARC. The contrast functions (CTFs) were determined using patch CTF and an initial stack of particles were picked using Blob picker followed by template picker after initial reconstruction. 18265163 particles were extracted from micrographs and were subjected to several rounds of 2D classifications followed by ab-initio reconstructions with subsequent heterogenous refinements in CryoSPARC that provided final set of 307,870 particles. Bayesian polished particles (Relion 4.0) resulted in reconstruction that was refined using non-uniform refinement function to 3.72 Å, with significant preferred orientation (cFAR = 0.01; SCF = 0.42). The resolution metrics provided here follow the gold-standard Fourier shell correlation 0.5 criterion (we adhered to a more stringent FSC criterion when determining the reported resolution due to preferred orientation). For illustration purposes and to aid model building, the cryo-EM map was processed with EMReady^72^ or blurred using gaussian filter in ChimeraX^73^. The individual component models from crystal structures were docked into the map using UCSF ChimeraX^73^ followed by flexible fitting with Namdinator^74^ with default parameters. The model was improved using iteratively Phenix real space refinement,^69^ with manual building in Coot^70^.

### CryoEM analysis of TGF-β3WD:zfBGo:TGFBR1:TGFBR2 complex and model refinement

Aliquots of 3 μL of each complex were applied to glow discharged UltrAuFoil r1.2/1.3 grids. The grids were blotted for 3 seconds at 100% humidity with force “2” and plunge frozen into liquid ethane using a Vitrobot Mark IV (Thermo Fisher). 7376 movies were recorded on a 300 keV Titan Krios microscope (Thermo Fisher) equipped with a FalconIV camera and a Selectris energy filter using 0.83 Å/pix. The total dose was set to 50 electrons per Å^2^. Movies were corrected using patch motion correction in CryoSPARC. The contrast functions (CTFs) of the flattened micrographs were determined using patch CTF and an initial stack of particles were picked using Blob picker followed by template picker after initial reconstruction. 3 x 10^6^ Particles were extracted from micrographs and were subjected to several rounds of 2D classifications followed by ab-initio reconstructions with subsequent heterogenous refinements in CryoSPARC that provided final set of 281,881 particles. Bayesian polished particles (Relion 4.0) resulted in reconstruction that was refined using non-uniform refinement function to 4.10 Å, with significant preferred orientation (cFAR = 0.05; SCF = 0.89). The resolution metrics provided here follow the gold-standard Fourier shell correlation 0.5 criterion (we adhered to a more stringent FSC criterion when determining the reported resolution due to preferred orientation). For illustration purposes and to aid model building, the cryo-EM map was processed with EMReady^72^. The crystal structure of the complex ^75^ was docked into the map using UCSF ChimeraX^73^ followed by flexible fitting with Namdinator^74^ with default parameters. The model was improved using iteratively using Phenix real space refinement^69^, with manual building in Coot^70^.

### CryoEM analysis of TGF-β1:ratBG_O_:(TGFBR2)_2_ complex and model refinement

Aliquots of 3 μL of complex sample was applied on UltrAuFoil r1.2/1.3 Grids covered with Graphene Oxide as described before^76^ and the grids were blotted for 3.5 s at force -1 at 4°C and plunge frozen into liquid ethane using a Vitrobot Mark IV (Thermo Fisher). 26,386 Movies were recorded on a 300 keV Titan Krios microscope (Thermo Fisher) equipped with a FalconIV camera and a Selectris energy filter using 0.921 Å/pix. The total dose was set to 50 electrons per Å^2^. Movies were corrected using motion correction as implemented incorporated into Relion 4.0 software package. The contrast functions (CTFs) were determined using GCTF and an initial stack of particles were picked using crYOLO with general model^77^. 1,016,267 particles extracted from micrographs using Relion 4.0 were imported into CryoSPARC and subjected to 3 rounds of initial 2D classification and several rounds of ab-initio reconstructions with subsequent heterogenous refinements that provided 101180 set of particles that were used to train TOPAZ and subsequent particle picking. 3978615 picked particles were extracted from micrographs and were subjected to several rounds of 2D classifications followed by ab-initio reconstructions with subsequent hetero refinements in CryoSPARC that provided final set of 230,626 particles. Bayesian polished particles resulted in reconstruction that was refined using non-uniform refinement function to an overall resolution 3.48 Å. The resolution metrics provided here follow the gold-standard Fourier shell correlation 0.143 criterion. For illustration purposes and to aid model building, the cryo-EM map was processed with EMReady^72^. The individual component models from AlphaFold2^75^ were docked into the map using UCSF ChimeraX ^73^ followed by flexible fitting with Namdinator^74^ with default parameters. The model was improved using iteratively Phenix real space refinement^69^, with manual building in Coot^70^ .

### CryoEM analysis of TGF-β:BG complex

Aliquots of 3 μL of complex sample was applied on UltrAuFoil r1.2/1.3. The grids were blotted for 3 seconds at 100% humidity with force “2” and plunge frozen into liquid ethane using a Vitrobot Mark IV (Thermo Fisher). 9496 Movies were recorded on a 300 keV Titan Krios microscope (Thermo Fisher) equipped with a FalconIV camera and a Selectris energy filter using 0.83 Å/pix. (including 2181 movies recorded on 20° tilted stage) The total dose was set to 55 electrons per Å^2^. Movies were corrected using patch motion correction in CryoSPARC. The contrast functions (CTFs) were determined using patch CTF and an initial stack of particles were picked using Blob picker followed by template picker after initial reconstruction. 8179080 Particles were extracted from micrographs and were subjected to several rounds of 2D classifications followed by ab-initio reconstructions with subsequent heterogenous refinements in CryoSPARC that provided final set of 256,006 particles. Final reconstruction that was refined using non-uniform refinement function yielded map at resolution 6.39Å. The resolution metrics provided here follow the gold-standard Fourier shell correlation 0.5 criterion (we adhered to a more stringent FSC criterion when determining the reported resolution due to preferred orientation).The individual component models from above structures or AlphaFold^75^ were docked into the map using UCSF ChimeraX^73^.

### Coupling of O6-(4-Aminomethyl-benzyl)guanine (BG-NH2) to CF640R-NHS

Reaction vial was charged with CF640R-NHS (MW: 929.19, 1 µmol, 1 mg) dissolved in 300 µl of DMSO and BG-NH2 (MW: 270.3, 1.4 µmol, 0.4 mg) dissolved in 200 µl DMSO. 3.0-fold molar excess of dry triethylamine (MW: 101.19, density: 0.726g/ml, 0.326 mg ∼0.44 µl) was added to reaction vial. The reaction was stirred under argon overnight at 30°C. The reaction was monitored using LC-MS (Cal M.W: 1084.49, Found: 1084.82) and was directly purified on PerkinElmer HPLC system using Solvent A (0.08%TFA and 0.1% acetonitrile in water) and Solvent B (0.1% TFA in acetonitrile) as an eluent in gradient 0 - 40% over 40 min to give the conjugated compound.

### Establishment of a stable HEK293T SNAP-BG cell line

To establish a stable HEK293T cell line expressing the SNAP-BG fusion protein, the pCDNA3.1+ plasmid containing the SNAP-BG fusion gene was linearized using the FspI restriction enzyme. The linearized plasmid was then transfected into HEK293 cells at 80% confluency using FuGENE HD reagent following the manufacturer’s protocol (Promega, Madison, WI, USA). Following transfection, positive selection was initiated by the addition of hygromycin at a concentration of 400 µg/ml. Cells were cultured until single colonies were visible and displayed 50% confluency.

### Evaluation of SNAP-BG expression, localization and activity

To assess SNAP-BG expression and correct subcellular localization, cells were grown on glass coverslips until reaching approximately 70% confluency. Subsequently, the cells were incubated with 30 nM SNAP-CF640R ligand for 20 minutes. After ligand incubation, excess dye was removed by washing three times with phosphate-buffered saline (PBS) and the cells were fixed with 4% paraformaldehyde (PFA), stained with DAPI and washed again 2X with PBS. Coverslips with fixed cells were mounted onto glass slides using mounting medium (ProLong^TM^ antifade reagent) and subjected to confocal microscopy analysis using a Leica SP8 confocal microscope.

The activity of SNAP-BG was assessed as reported before^20^. Briefly, HEK293 cells expressing SNAP-BG were treated with low concentration (20, 40 pM) of TGF-β2 for 15 min. Cells were then washed, lysed using lysis buffer and assayed for the presence of pSMAD2 using Western Blot. Presence of SNAP-BG was also confirmed by probing the membrane with anti-SNAP-tag antibody.

### Fluorescent labeling of TGF-β2 and InhA

Purified TGF-β2 (0.5 mg) or InhA (0.1 mg) at a concentration of 2 mg mL^-1^ in 10 mM acetic acid was combined with 100 µL of phosphate-buffered saline (PBS) pH 7.2 and 60 µL of dimethyl sulfoxide (DMSO). The pH was adjusted to approximately 7.0. A 5:1 molar excess of the succinimide ester of CF640R fluorescent dye (Biotium) or Alexa Fluor^TM^ 647 was then added to the protein solution. The labeling reaction was allowed to proceed for 60 min and was quenched by adding 100 µL of 100 mM Tris-HCl pH 8.0. The labeled protein was then dialyzed against 10 mM acetic acid and purified using a cation exchange MonoS column (Cytiva) with a gradient of NaCl. Fractions containing the labeled protein were identified using gel electrophoresis and the ImageQuant™ 800 fluorescent gel documentation system (Cytiva). Pooled fractions containing the labeled protein were dialyzed against 50 mM acetic acid, concentrated, and stored at -20°C until further use.

### Competition binding assay

HEK293T cells stably expressing either the SNAP-BG construct or a control strain were seeded onto 18 mm coverslips placed in 12-well plates at a density of 10^5^ cells per well. Upon reaching approximately 80% confluency, cells were treated with 20 ng mL^-1^ of fluorescently labelled TGF-β2 or 20 ng mL^-1^ InhA. After 10 minutes, a 100-fold excess of the indicated unlabeled ligand was added and cells were further incubated for 15 min. Subsequently, SNAP-CF568 at final concentration of 30 nM was introduced, and cells were incubated for an additional 15 min. Excess dye was removed by washing the cells three times with phosphate-buffered saline (PBS). Cells were then fixed with 4% paraformaldehyde (PFA), stained with DAPI and washed again 2X with PBS. Fixed cells on coverslips were carefully mounted onto glass slides using mounting medium (ProLong^TM^ antifade reagent). Subsequently, the samples were subjected to detailed confocal microscopy analysis using a Leica SP8 confocal microscope.

### Confocal microscopy and image processing

The samples were subjected to confocal microscopy analysis using a Leica SP8 confocal microscope using an HC PL APO CS2 20x/ 0.75 IMM objective at 25 °C with the following confocal settings: pinhole 1 airy unit, scan speed 400 Hz unidirectional, format 1024 x 1024 pixels at 16 bit. Images were collected using hybrid detectors and an argon, 561 nm and 633 nm lasers with 2x line averaging. For each field of view (FOV), a stack of 25 images was collected with a z-step of 0.5 µm. Collected images were processed using Fiji software ^78^. Images from the stack were combined into a single z-projection, and a threshold was established using the Otsu method for automatic thresholding, using the SNAP-BG signal channel as source. The intensity of the defined areas was measured for both the SNAP-BG and TGF-β2/inhibinA channels. The ratio of intensities for each channel was calculated and plotted for visualization using scatter plots in GraphPad Prism version 10.1.1 for Mac, GraphPad Software, Boston, Massachusetts USA.

### Data Availability

This paper does not report any original code. Any additional information required to reanalyze the data reported in this paper is available from the lead contacts upon request. Structural data has been deposited to Protein Data Bank (PDB) or/and Electron Microscopy Data Bank (EMDB) and will be released upon publication or can be made available by contacting the corresponding author. Plasmids generated in this study are maintained in the laboratories of Andrew Hinck and Caroline Hill and will be made available upon request.

---

**Extended Data Figure 1.**
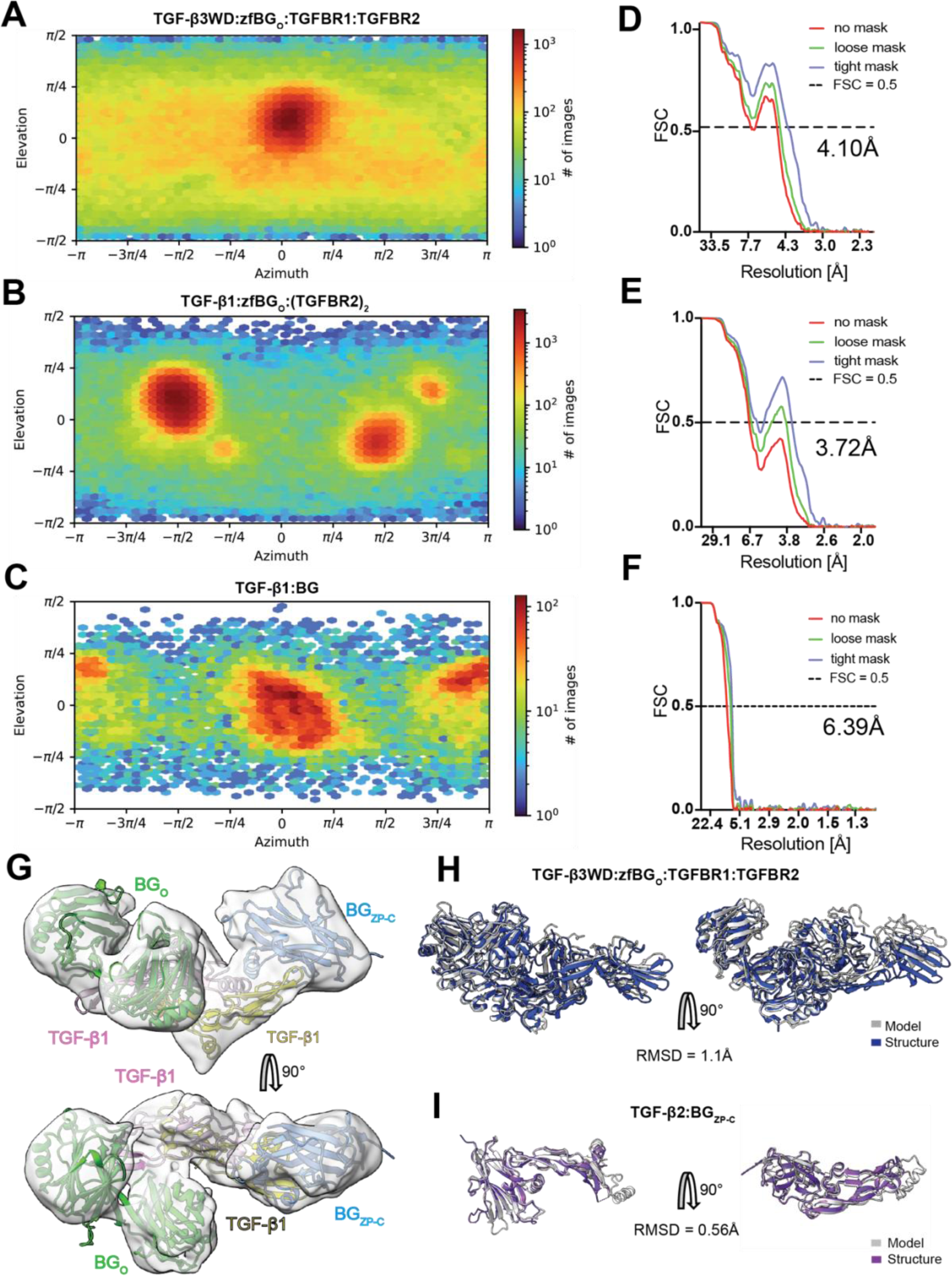
Structural details of the Cryo-EM complexes and AF2M models. (A–F) Particle view distributions for presented 3D reconstructions of TGF- β3WD:zfBG_O_:TGFBR1:TGFBR2 (A), TGF-β1:zfBG_O_:(TGFBR2)_2_ (B), or TGF-β1:BG (C), together with Fourier Shell Correlation (FSC) curves (D, E, F, respectively). (G) Low resolution Cryo-EM map of the TGF-β:BG complex with fitted model to the density. (H, I) AF2M models of TGF-β3WD:BG_O_:TGFBR1:TGFBR2 (H) and TGF-β2:BG_ZP-C_ (I). The complexes are overlaid with crystal structures of TGF- β3WD:zfBG_O_:TGFBR1:TGFBR2 and mmTGF-β2:BG_ZP-C_, respectively.

**Extended Data Figure 2:**
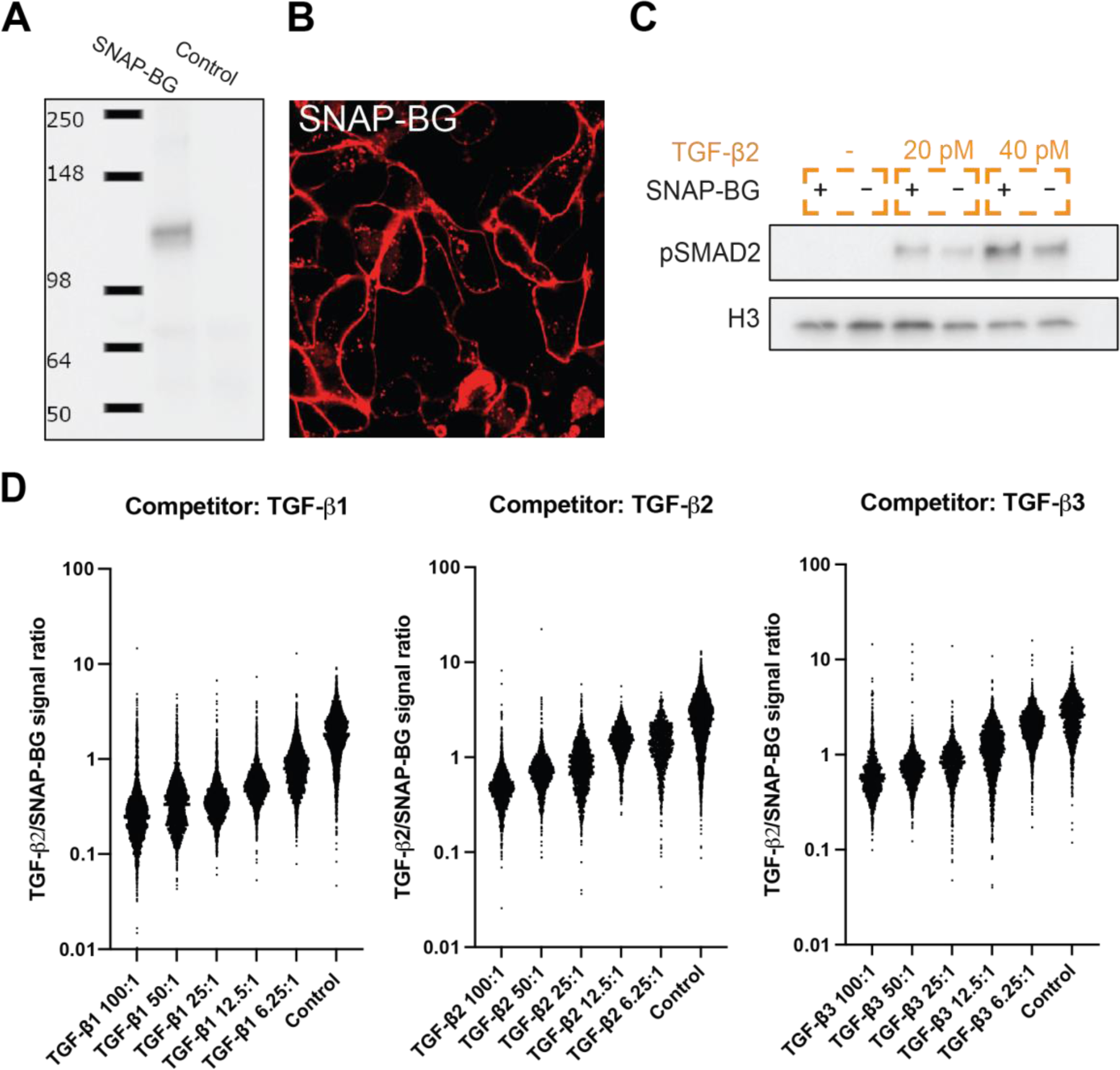
Characterization of SNAP-Betaglycan (SNAP-BG) in HEK293T Cells. (A) SNAP-BG expression in HEK293 cells. Western blot analysis of SNAP-BG expression in stably transfected HEK293T cells with pCDNA3.1 plasmid harbouring SNAP-BG. Proteins were resolved on SDS gel and probed with an antibody specific to the SNAP-tag. A distinct band is observed at approximately 120 kDa, consistent with the expected molecular weight of the ΔGAG SNAP-BG fusion protein. (B) SNAP-BG has membrane localization. Confocal microscopy images showing SNAP-BG conjugated with a fluorescent ligand (shown in red), predominantly localized on the cell membrane. (C) Western blot analysis demonstrates a dose-dependent increase in phosphorylated SMAD2 (pSMAD2) levels in the presence of SNAP-BG. (D) Concentration-dependent effects of TGF-β isoforms as competitors in the binding assay. As the concentration of each TGF-β isoform increases (TGF-β1, TGF-β2, TGF-β3), there is a corresponding decrease in the colocalization ratio of TGF-β2- CF640R/SNAP-BG, indicating assay sensitivity to ligand concentration.

**Extended Data Figure 3.**
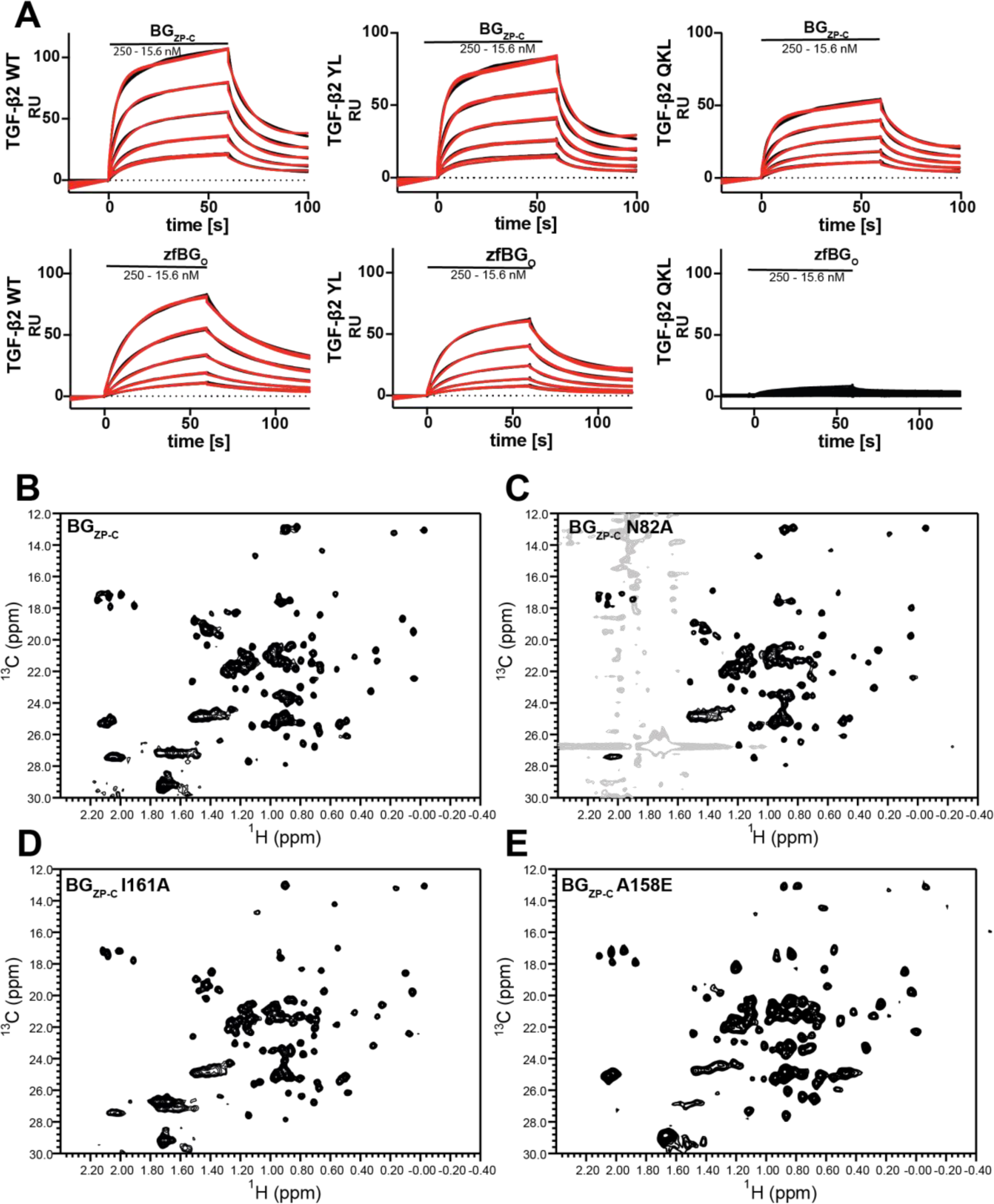
Mutations affecting TGF-β2 binding cause only local structural changes. (A) Sensorgrams from SPR experiment where zfBG_O_ WT or BG_ZP-C_ WT were injected over immobilized TGF-β2 WT, TGF-β2 (Q57A, K60A, L64A located on heel helix), or TGF-β2 (Y50A, L51A located in pre-helix extension). zfBG_O_ binding was severely affected only by mutations in the heel helix, while the binding of BG_ZP-C_ was mostly unaffected. (B–E) Natural abundance methyl ^1^H-^13^C correlation NMR spectra indicating minimal perturbations in the chemical environment of binding affecting mutants of BG_ZP-C_, reassuring that the mutations lead to localized structural alterations rather than a widespread change in conformation.

**Extended Data Figure 4.**
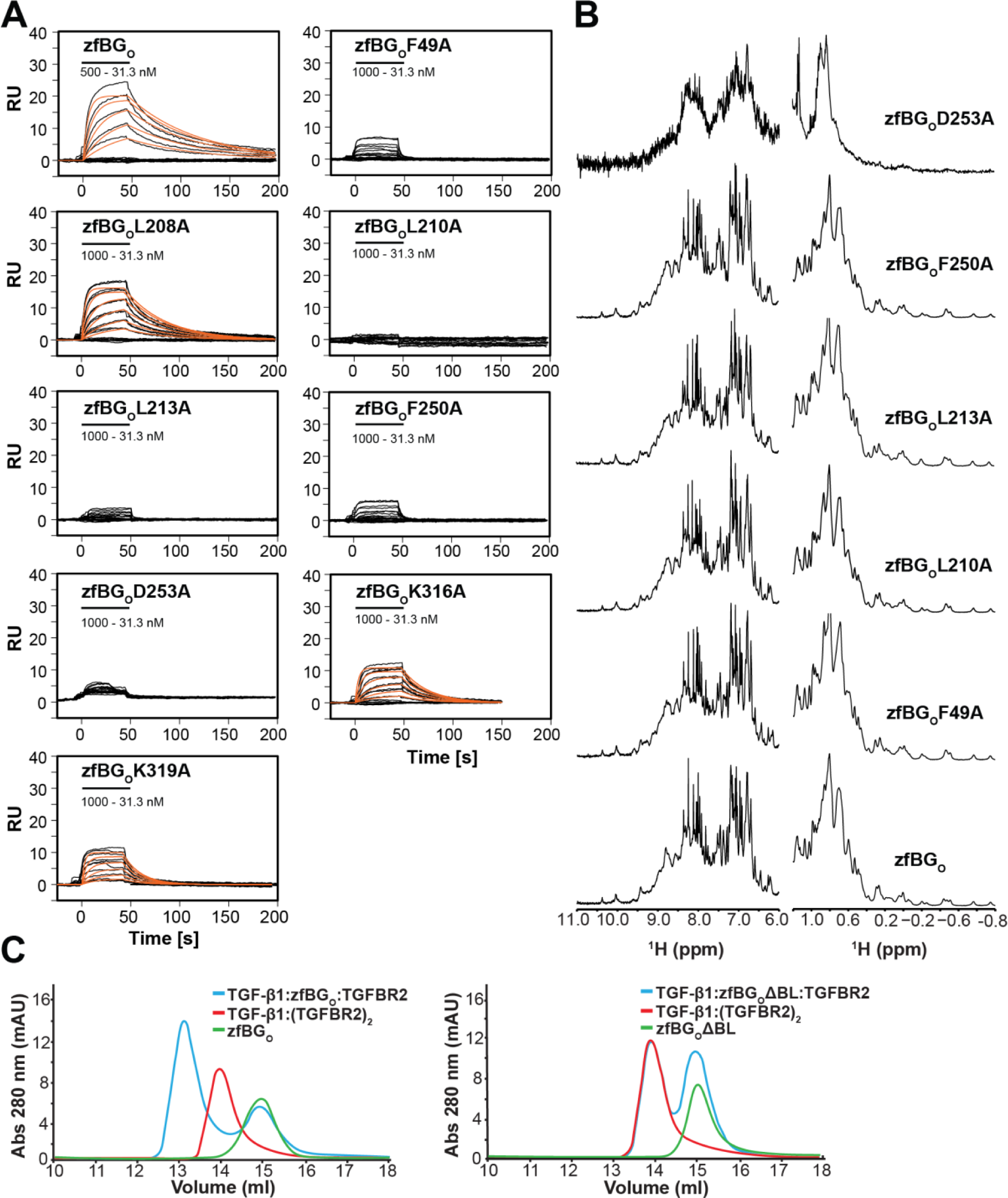
zfBG_O_ mutants within TGF-β:zfBG_O_ binding interface. (A) SPR sensorgrams of zfBG_O_ mutants binding to immobilized TGF-β2. (B) 1D proton NMR spectra, covering both amide and methyl spectral regions. The spectra underscore that mutations predominantly result in localized alterations to the binding interface, without extensive perturbation of the overall protein structure. Notably, the D253A mutant exhibits signal broadening across the spectra, hinting at more widespread structural changes that may suggest alterations in protein folding. (C) SEC profiles of TGF-β1:zfBG_O_:(TGFBR2)_2_ and TGF-β1:(TGFBR2)_2_ complexes (left panel) or TGF-β1:zfBG_O_ΔBL:(TGFBR2)_2_ and TGF-β1:(TGFBR2)_2_ complexes (right panel), where zfBG_O_ΔBL corresponds to zfBG_O_ with BL fully replaced with unrelated sequence. No shift in the TGF-β1:zfBG_O_ΔBL:(TGFBR2)_2_ elution volume together with the presence of second peak eluted at the BG_O_ volume is indicative of zfBG_O_ΔBL not forming a complex with TGF-β1:(TGFBR2)_2_.

**Extended Data Figure 5.**
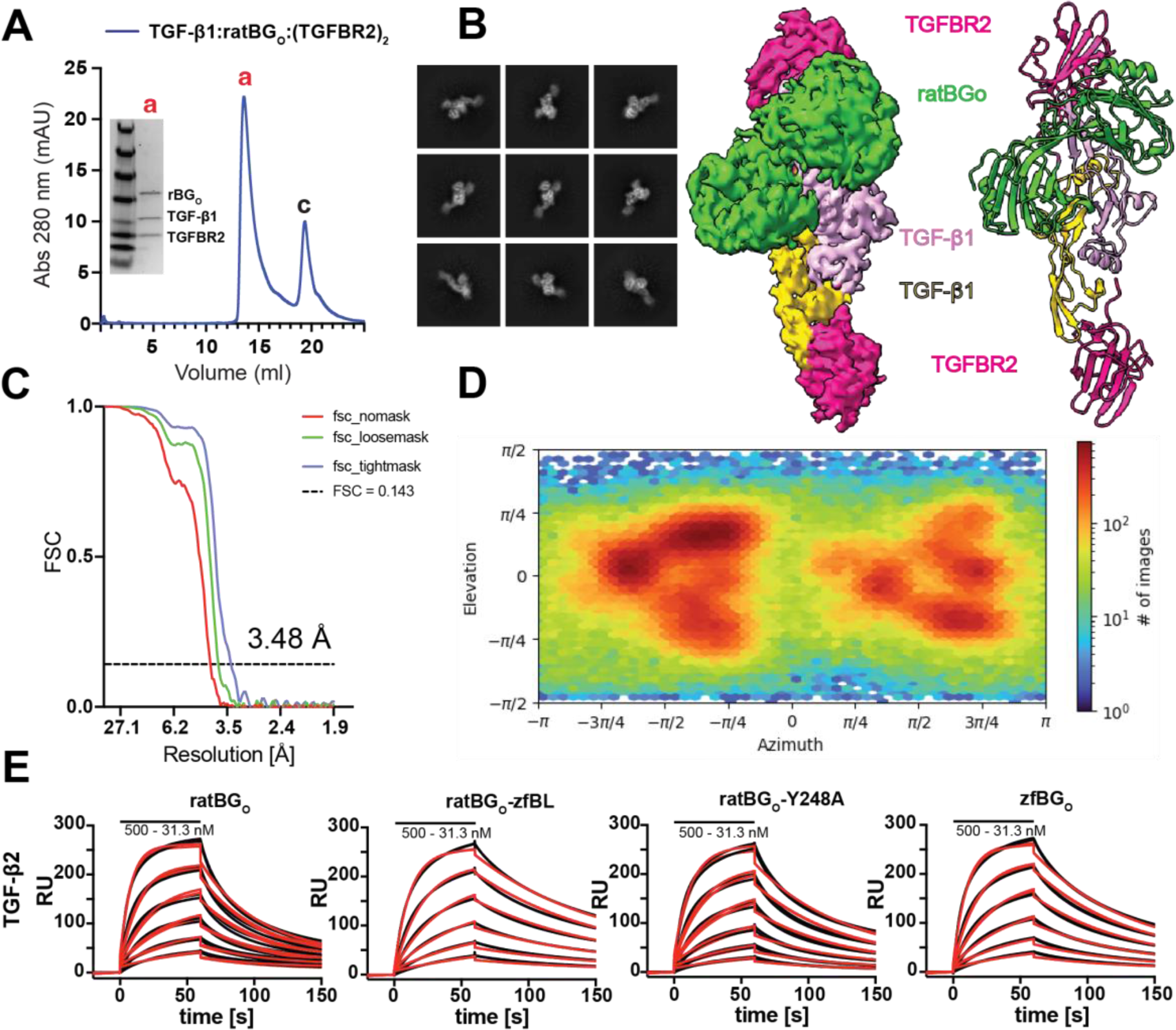
SPR validation of ratBG_O_ mutants binding to TGF-β and structural details of the CryoEM. (A) SEC purified, assembled complex of TGF-β1:ratBG_O_:(TGFBR2)_2_. (B) Selected 2D classes and Cryo-EM map with a model built into the density (C) FSC curve of the final map refinement with particle view distribution (E). (D) SPR sensorgrams showing binding of ratBG_O_, zfBG_O_, as well as two mutants, ratBG_O_zfBL and ratBG_O_ –Y248A, to TGF-β2.

**Extended Data Figure 6.**
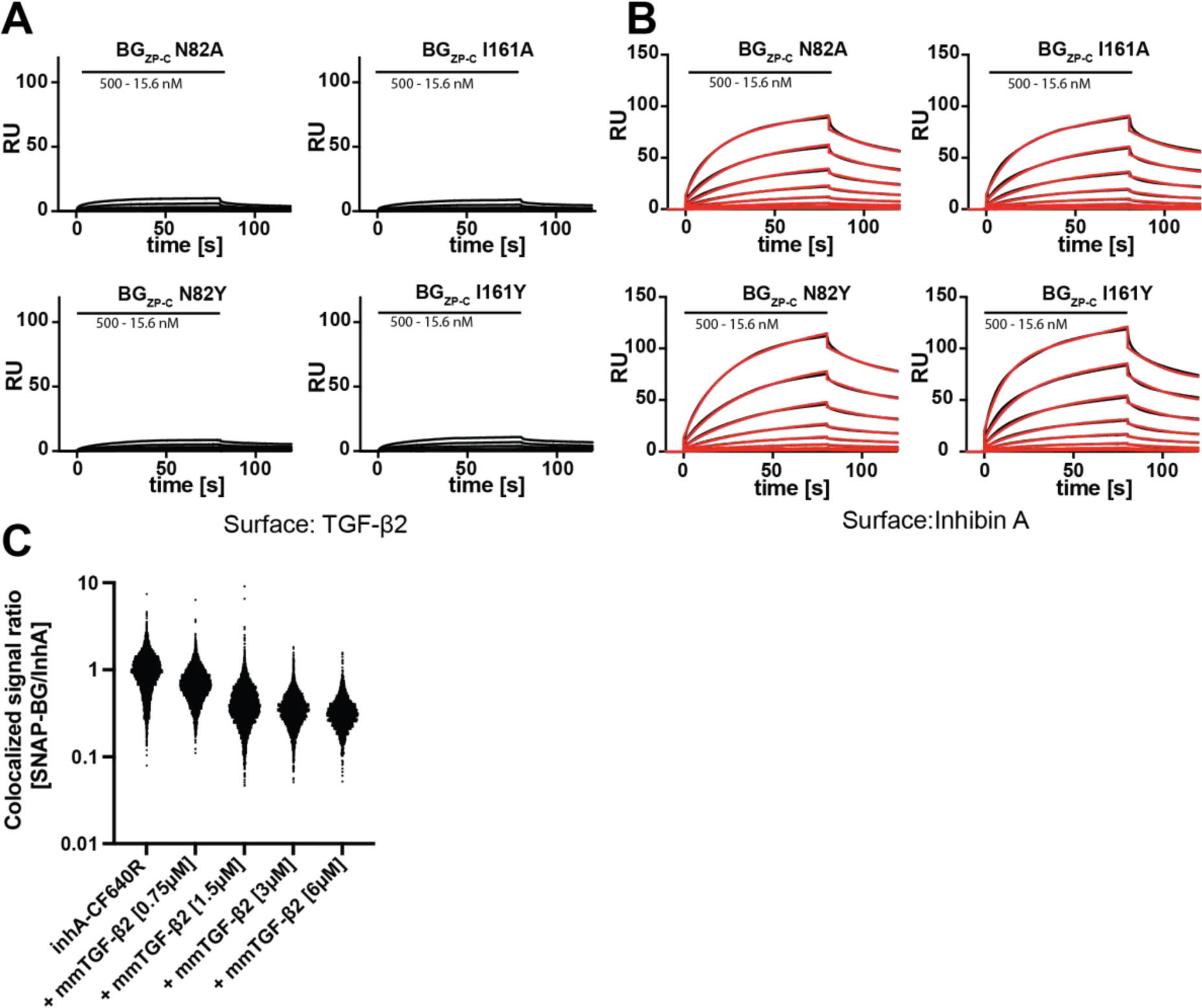
InhA uses different, but overlapping to TGF-βs interface to bind to BG_ZP-C_. (A, B) SPR sensograms of selected mutants of BG_ZP-C_ binding to immobilized TGF- β2 (A) or InhA (B). (C) Quantification of concentration-dependent effect of mmTGF-β2 on InhA-CF640R signal strength.

**Suplementary Table 1.**
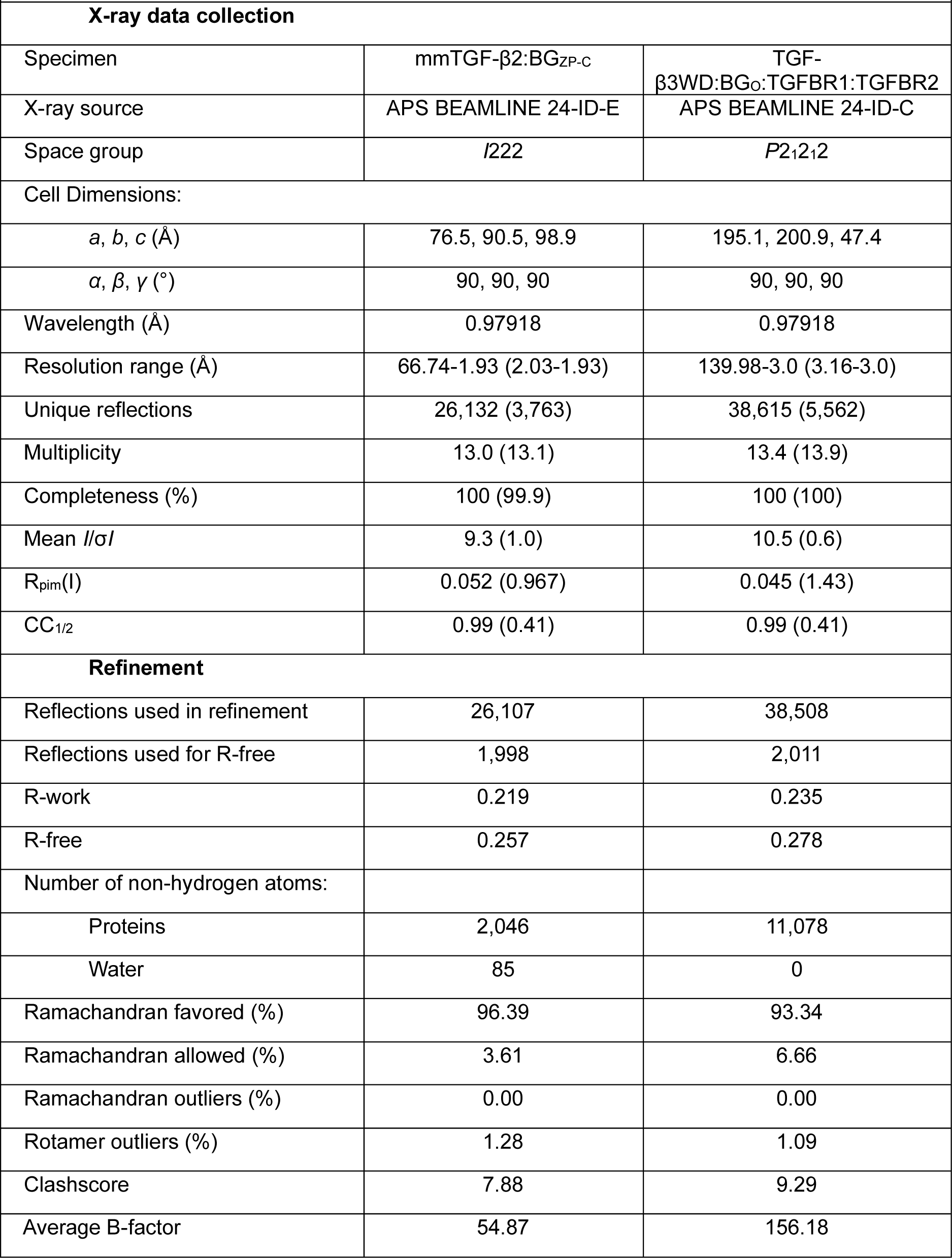
X-ray data collection and refinement parameters. Values in parentheses represent the highest resolution shell.

**Suplementary Table 2.** CryoEM data collection and refinement parameters.

| **Supplementary Table 2.** CryoEM data collection and refinement parameters |  |  |  |  |
| --- | --- | --- | --- | --- |
| Specimen | -TGF-β3WD<br>-zfBG<sub>o</sub><br>-TGfBR1<br>-TGfBR2 | -TGF-β1<br>-zfBG<sub>o</sub><br>-(TGfBR2)<sub>2</sub> | -TGF-β1<br>-BG | -TGF-β1<br>-ratBG<sub>o</sub><br>-(TGfBR2)<sub>2</sub> |
| **Data collection** |  |  |  |  |
| Magnification | 168675 x | 194444 x | 168675 x | 152009 x |
| Voltage | 300 | 300 | 300 | 300 |
| Electron exposure (e-/Å<sup>2</sup>) | 50 | 50 | 55 | 50 |
| Defocus range (μm) | -0.75 to -2.25 | -0.75 to -2.25 | -0.75 to -2.25 | -1.5 to -3.5 |
| Pixel size | 0.83 | 0.72 | 0.83 | 0.921 |
| Symmetry imposed | C1 | C1 | C1 | C1 |
| Initial particle images | 3 001 720 | 11 800 000 | 6 500 000 | 3 978 615 |
| Final particle images | 281 881 | 307 870 | 258 006 | 235 089 |
| Map resolution (Å) | 4.10 | 3.72 | 6.39 | 3.48 |
| cFAR | 0.05 | 0.01 | 0.36 | 0.4 |
| SCF | 0.89 | 0.42 | 0.97 | 0.80 |
| FSC threshold | 0.5 | 0.5 | 0.5 | 0.143 |
| **Refinement** |  |  |  |  |
| Map sharpening B factor (Å<sup>2</sup>) | 99.9 | 63.5 | 457 | 172.1 |
| Chains | 5 | 5 | 3 | 5 |
| Non-hydrogen atoms | 5766 | 6093 | NA | 6120 |
| Protein residues | 735 | 769 | NA | 770 |
| Ligands | 0 | 0 | 0 | 0 |
| **Validation** |  |  |  |  |
| - MolProbity score | 2.03 | 1.86 | NA | 2.13 |
| - Clashcore | 7.29 | 7.06 | NA | 9.26 |
| - Poor rotamers (%) | 2.25% | 1.86% | NA | 2.56% |
| **Ramachandran plot** |  |  |  |  |
| - Favored | 94.76% | 96.05% | NA | 95.39% |
| - Allowed | 5.24% | 3.95% | NA | 4.61% |
| - Disallowed | 0.00% | 0.00% | NA | 0.00% |

**Suplementary Table 3.**
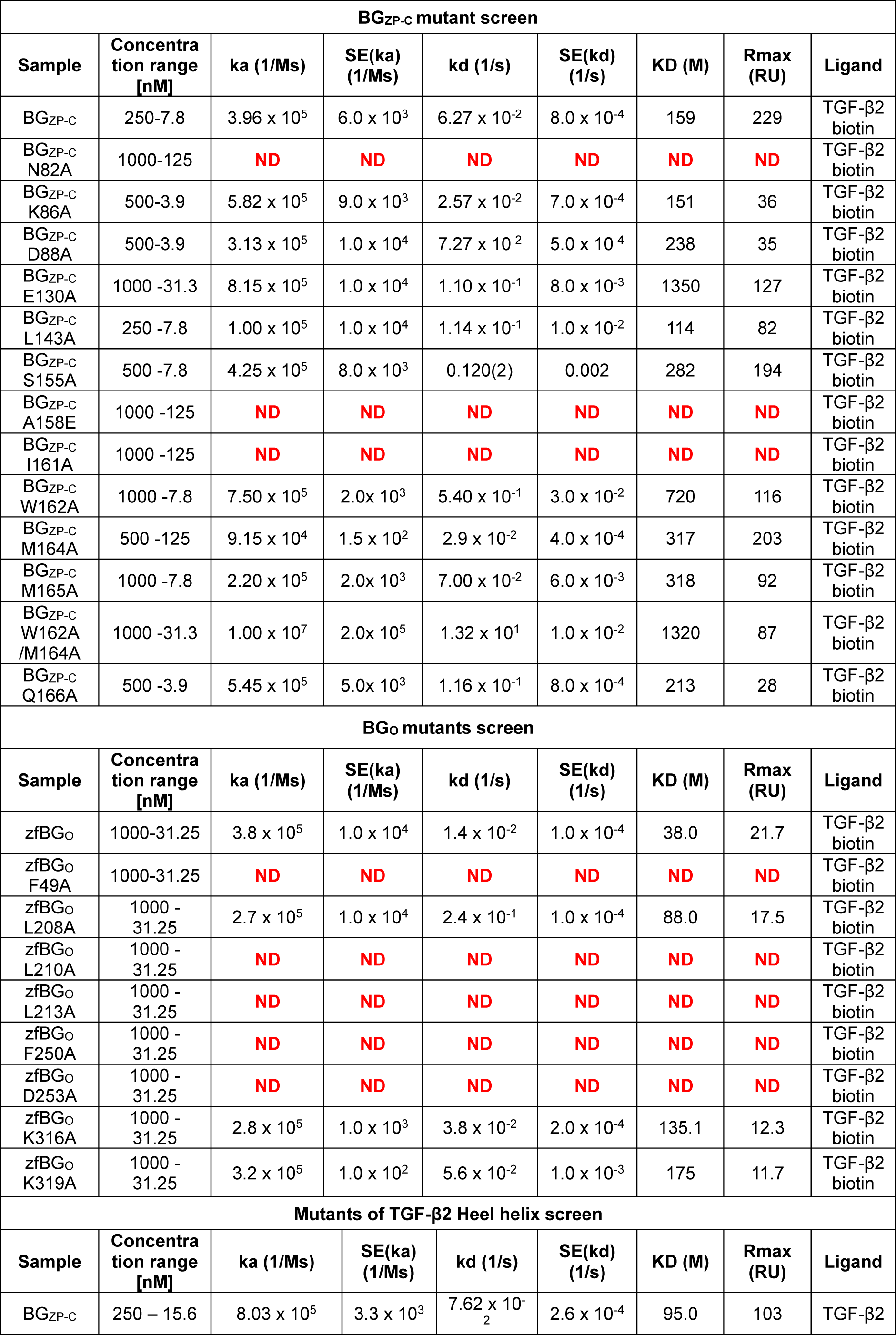
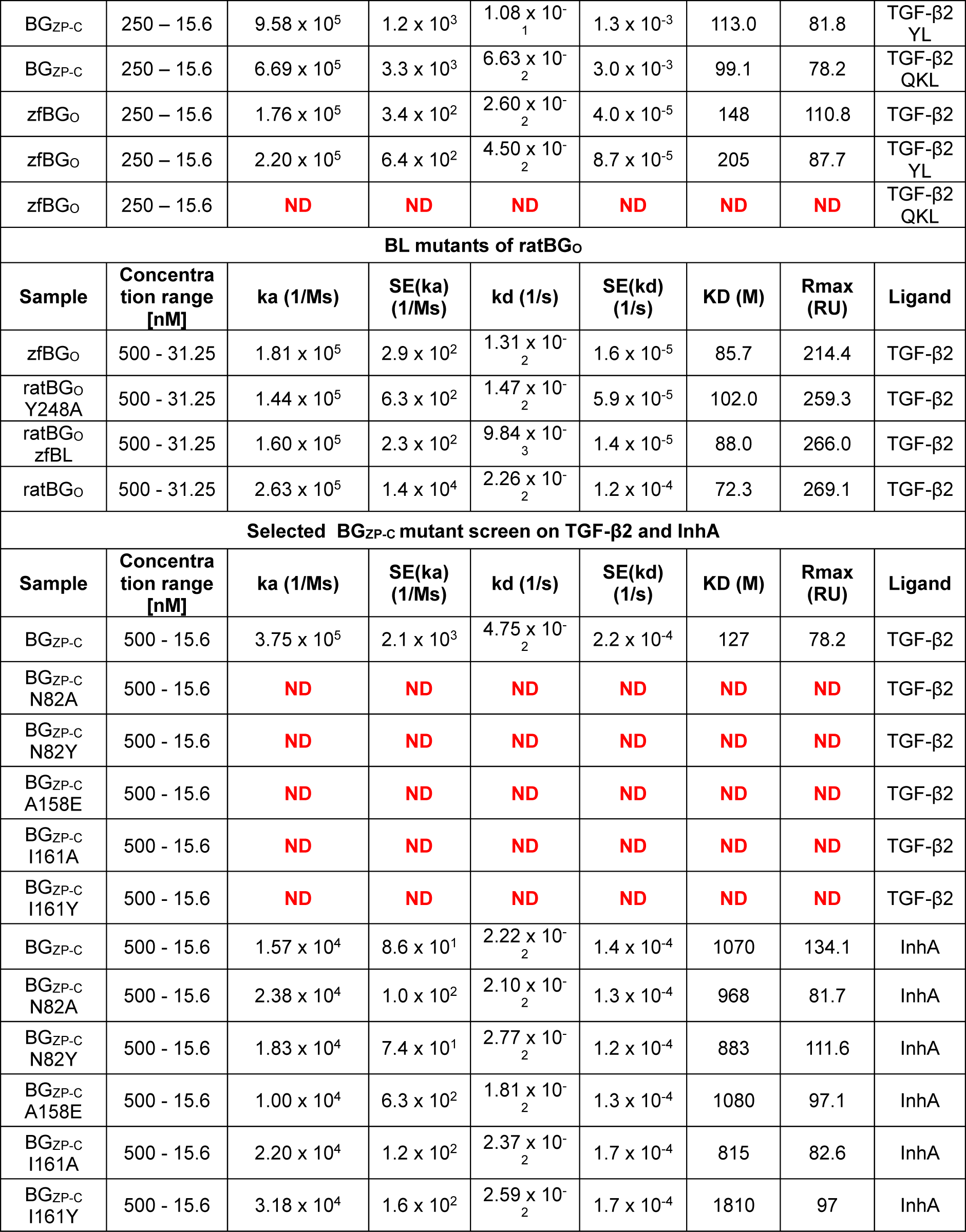
Parameters of SPR binding measurements.

